# Mechanistic Insights into Crosstalk of Tet(X) and MCR-1, Two Resistance Enzymes Co-produced by A Single Plasmid

**DOI:** 10.1101/2020.03.06.981738

**Authors:** Yongchang Xu, Lizhang Liu, Huimin Zhang, Youjun Feng

## Abstract

Tigecycline and colistin are few of last-resort defenses used in anti-infection therapies against carbapenem-resistant bacterial pathogens. The successive emergence of plasmid-borne *tet*(X) tigecycline resistance mechanism and mobile colistin resistance (*mcr*) determinant, renders them clinically ineffective, posing a risky challenge to global public health. Here, we report that co-carriage of *tet*(X6) and *mcr-1* gives co-resistance to both classes of antibiotics by a single plasmid in *E. coli*. Genomic analysis suggested that transposal transfer of *mcr-1* proceeds into the plasmid pMS8345A, in which a new variant *tet*(X6) is neighbored with Class I integron. The structure-guided mutagenesis finely revealed the genetic determinants of Tet(X6) in the context of phenotypic tigecycline resistance. The combined evidence *in vitro* and *in vivo* demonstrated its enzymatic action of Tet(X6) in the destruction of tigecycline. The presence of Tet(X6) (and/or MCR-1) robustly prevents the accumulation of reactive oxygen species (ROS) induced by tigecycline (and/or colistin). Unlike that *mcr-1* exerts fitness cost in *E. coli, tet*(X6) does not. In the *tet*(X6)-positive strain that co-harbors *mcr-1*, tigecycline resistance is independently of colistin resistance caused by MCR-1-mediated lipid A remodeling, and vice versa. Co-production of Tet(X6) and MCR-1 gives no synergistic delayed growth of the recipient *E. coli*. Similar to that MCR-1 behaves in the infection model of *G. mellonella*, Tet(X6) renders the treatment of tigecycline ineffective. Therefore, co-transfer of such two AMR genes is of great concern in the context of “one health” comprising environmental/animal/human sectors, and heightened efforts are required to monitor its dissemination.

**Author summary:** We report that *tet*(X6), a new tigecycline resistance gene, is co-carried with the other resistance gene *mcr-1* by a single plasmid. Not only have we finely mapped genetic determinants of *tet*(X6), but also revealed its biochemical action of tigecycline destruction. Crosstalk of Tet(X6) with MCR-1 is addressed. Tet(X6) tigecycline resistance is independently of MCR-1 colistin resistance, and vice versa. Similar to MCR-1 that renders colistin clinically ineffective, Tet(X6) leads to the failure of tigecycline treatment in the infection model of *G. mellonella*. This study extends mechanistic understanding mechanism and interplay of Tet(X6) and MCR-1, coproduced by a single plasmid. It also heightens the need to prevent rapid and large-scaled spread of AMR.

## Introduction

Antimicrobial resistance is an increasingly-devastating challenge in the context of “one health” that covers the environmental, animals and human sectors. Colistin is one of cationic antimicrobial polypeptides (CAMP) with an initial target of the surface-anchored lipid A moieties on the Gram-negative bacterium ^1^. In contrast, tigecycline is the third-generation of tetracycline-type antibiotic, which interferes the machinery of protein synthesis of both Gram-negative, and Gram-positive bacteria ^2^. In general, both colistin and tigecycline are an ultimate line of antibiotics to combat against lethal infections with carbapenem-resistant pathogens ^3,4^. Unfortunately, the emergence and global distribution of MCR family of mobile colistin resistance (*mcr-1* ^5,6^ to *mcr-10* ^7^) has potentially threatened the renewed interest of colistin in clinical therapies ^8^. The majority of transferable colistin resistance depends on the surface lipid A remodeling by the MCR enzymes via the “ping-pong” trade-off ^9^.

In addition to the two well-known actions, efflux and ribosome protection ^10,11^, antibiotic degradation also constitutes in part the mechanism of tigecycline resistance ^12-14^. The Tet(X) enzyme is a class of flavin-requiring monooxygenase ^15,16^, which possesses the ability of modifying tetracycline and its derivatives (like the glycylcycline, tigecycline) ^13^. In general, Tet(X) inactivates tigecycline to give 11a-Hydroxytigecycline, rendering the carrier host bacterium insusceptible to tigecycline ^13^. In particular, functional meta-genomics of soils performed by Forsburg *et al*. ^17^ revealed a number of new tetracycline destructases (10 in total, namely Tet47 to Tet56). Among them, *tet56* is only one exclusively from the human pathogen *Legionella longbeachae*, indicating its potential spread from environments to clinical sector ^17^. Subsequent structural studies suggested that these Tet tetracycline destructases accommodate antibiotics in diverse orientation, which highlights their architectural plasticity ^12^. Indeed, the discovery of an inhibitor blocking the entry of flavin adenine dinucleotide cofactor into Tet(50) enzyme also paved a new way to reversing tigecycline resistance ^12^.

Very recently, two new variants [*tet*(X4) and *tet*(X5)] of *tet*(X)-type determinants that encode a tigecycline-inactivating enzyme, was found to spread by distinct plasmids of *Escherichia coli* in China ^18,19^. Although the limited distribution of *tet*(X4) thus far ^20^, it constitutes an expanding family of Tet(X) resistance enzyme [Tet(X) ^21,22^ to Tet(X5) ^19^], and raises the possibility of rendering tigecycline (and even the newly-FDA approved eravacycline, the fourth-generation of tetracycline derivatives ^23^) clinically ineffective. Worrisomely, the co-transfer of *mcr-1* and *tet*(X) probably promotes the emergence of a deadly superbug with the co-resistance to polymyxin and tigecycline. However, this requires further epidemiological evidence.

Here, we report that this is the case. It underscores an urgent need to monitor and evaluate a potential risk for the convergence of Tet(X) tigecycline resistance to MCR colistin resistance by a single highly-transmissible plasmid in an epidemic ST95 lineage of virulent *E. coli* ^24^.

## Results and Discussion

### Discovery of *tet*(X6), a new variant of Tet(X) resistance enzyme

To address this hypothesis, we systematically screened the whole NCBI nucleotide database, in which each of the six known *tet*(X) variants [*tet*(X) to *tet*(X5)] acts as a query. Among them, it returned six hits with the significant score (97%-99% identity) when we used the *tet*(X3) of *Pseudomonas aeruginosa* (1137bp, Acc. no.: AB097942) as a searching probe. The resultant hits corresponded to four contigs of uncultured bacterium and two plasmids. These contigs include TE_6F_Contig_7 (3328bp, Acc. no.: KU547125), TG_6F_Contig_3 (3323bp, Acc. no.: KU547130), TG_7F_Contig_3 (2575bp, Acc. no.: KU547185); and TE_7F_Contig_3 (2588bp, Acc. no.: KU547176). The matched plasmids refer to pMS8345A (241,162bp; Acc. no.: CP025402) ^24^ and p15C38-2 (150,745bp; Acc.no.: LC501585), in which the gene of MS8345_A00031 exhibits 97.1% identity and 100% coverage when compared to *tet*(X3) (**Fig. S1**). This *tet*(X3)-like gene, MS8345_A00031, is thereafter renamed *tet*(X6) (Acc. no.: BK011183) in this study (**Fig. 1A**). Strikingly, we found that this plasmid co-harbors the *mcr-1* gene encoding a <u>p</u>hospho<u>e</u>thanol<u>a</u>mine (PEA)-lipid A transferase ^25^. Because that the plasmid is detected in an epidemic clone of ST95 <u>Ex</u>traintestinal <u>P</u>athogenic *<u>E</u>. <u>c</u>oli* (ExPEC) ^24^, this single plasmid pMS8345A possesses the potential to confer its recipient host *E. coli* the co-resistance to both tigecycline and colistin. It is unusual, but not without any precedent. In fact, *mcr-1* has ever coexist with *bla*_NDM_ in a single isolate ^26^ and even a single plasmid ^27^.

**Figure 1.**
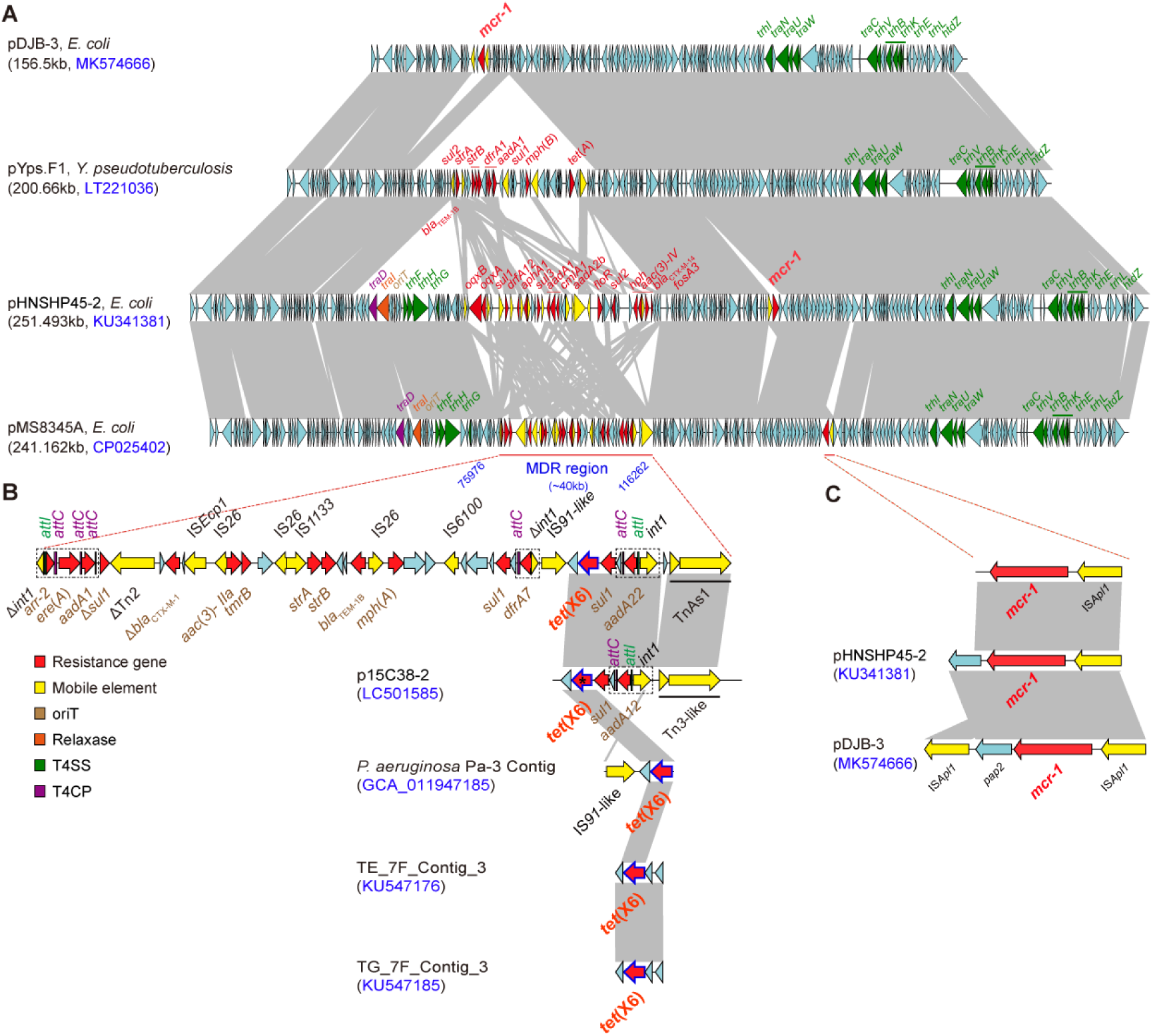
Genetic context-based analyses for co-transfer of *tet*(X6) and *mcr-1* by a single plasmid. **A.** Co-linear genome alignment of the *tet*(X6), and *mcr-1*-coharboring plasmid pMS8345A with other three IncHI2-type mega plasmids (pDJB-3, pYps.F1, and pHNSHP45-2) The backbones of the four resistance plasmids are of high similarity. Except that pYps.F1 (Acc. no.: LT221036) of *Yersinia pseudotuberculosis* lacks *mcr-1*, all the other three resistance plasmids here carry this gene. pDJB-3 (Acc. no.: MK574666) is a new variant of the *mcr-1*-containing plasmid we reported here. Unlike that pDJB-3 lacking multidrug resistance region (MDR), the remaining three plasmids possess different MDRs. ICEs are colored as follows: *oriT* in brown, the relaxase-encoding gene in orange, T4SS in green, and T4CP in purple. **B.** Fine dissection of the *tet*(X6)-positive MDR regions in pMS8345A and p15C38-2 As for this unique MDR region in pMS8345A, three Class 1 integrons are circled by dashed lines, 14 kinds of antibiotic resistance determinants like *addA1* (with red arrows), are integrated, and no less than 10 different mobile elements such as IS*Ecp1* and Tn*As1* (indicated with yellow arrows) are determined. In particular, a new *tet*(X) variant of tigecycline resistance enzymes, named *tet*(X6) (in bold red letters), is adjacent to *sul1* that conferring resistance to sulfonamide. Additionally, five *attL* sites (highlighted in purple) are detected. The gene environment of *tet*(X6) on pMS8345A is highly matched to the counterpart on plasmid p15C38-2, with an exception of a nucleotide deletion of “A” on the position of 287. This A287 deletion of *tet*(X6) highlighted with the symbol of “*” results in a frameshift and premature termination of its corresponding ORF (1137bp vs 282bp; 378aa vs 93aa). In contrast, it gives limited overlap with the three *tet*(X6)-containing contigs, *P. aeruginosa* Pa-3 contig, TE_7F_Contig_3, and TG_7F_Contig_3. **C.** Genetic context of the *mcr-1*-centering transposal region in pMS8345A, pHNSHP45-2, and pDJB-3 Colored arrows indicate ORFs and the shaded region depicts sequence similarity (70%-100%). The resistance genes are highlighted in red and mobile elements are highlighted in yellow. Designations: pDJB-3 (156.5kb, IncHI2/*E. coli*, China/2017, Acc. no.: MK574666); pYps.F1 (200.66kb, IncH1/*Y. pseudotuberculosis*, France/2017, Acc. no.: LT221036); pHNSHP45-2 (251.5kb, IncHI2/*E. coli*, China/2013, Acc. no.: KU341381); p15C38-2 (150.745kb, IncA/C2, *E. coli*/Japan/2003, Acc. no.: LC501585); *Pseudomonas aeruginosa* strain Pa-3 contig (3.796kb, *P.aeruginosa*, Pakistan/2016, Acc. no.: GCA_011947185.1); Uncultured bacterium clone TE_7F_Contig_3 (Acc. no.: KU547176); Uncultured bacterium clone TG_7F_Contig_3 (Acc. no.: KU547185) and pMS8345A (241.162kb, IncHI2/*E. coli*, Qutar/2017, Acc. no.: CP025402).

### Characterization of a *tet*(X6)-harboring plasmid

The *tet*(X6)-positive pMS8345A (Acc. no.: CP025402) that attracts us much attention during the period of *in silico* search, is an IncHI2-type large plasmid (∼241kb). In fact, it was initially discovered by Beatson and coworkers ^24^ from a virulent lineage of *E. coli* with <u>m</u>ultiple <u>d</u>rug <u>r</u>esistance (MDR) through the routine screen of colistin resistance (**Fig. 1A**). Generally, this plasmid has an average GC% value of 46.29%, and is predicted to contain 548 putative open reading frames (ORFs). Importantly, pMS8345A is found to possess a series of <u>I</u>ntegrative and <u>C</u>onjugative <u>E</u>lements (ICEs) using the web-based tool of oriTfinder (**Fig. 1A**) ^28^. Although that pMS8345A is not accessible in China right now, we applied its 2 surrogate plasmids, pHNSHP45 and pDJB-3 (**Fig. 1A**), in the conjugation assays. In general consistency with an observation of Zhi *et al*. ^29^, the efficiency of pHNSHP45-2 transfer is calculated to be 3.7×10^−5^ in our experiment of conjugations. In contrast, pDJB-3 can’t survive in the conjugation trials. Unlike pDJB-3 that carries T4SS alone (**Fig. 1A**), pHNSHP45-2 is fulfilled with all the four essential modules for self-transmission, namely *oriT* region, relaxase gene, type IV coupling protein (T4CP) gene and type IV secretion system (T4SS) (**Fig. 1B**). Notably, the aforementioned modules are shared by the plasmid pM8345A and pHNSHP45-2 (**Fig. 1B**). Taken together, we believed that pM8345A is self-transmissible.

The hallmark of pMS8345A lies in two disconnected/unique resistance regions, one of which refers to the *tet*(X6)-bearing MDR region of appropriate ∼40kb long (**Fig. 1B**), and the other denotes the *mcr-1*-containing region (**Fig. 1C**). In total, 11 kinds of antimicrobial resistance (AMR) genes have been recruited and integrated into this unusual MDR region (**Fig. 1B**), giving multiple drug resistance. Apart from tigecycline resistance caused by *tet*(X6), colistin resistance arises from the “*mcr-1*-IS*Apl1*” transposon alone (**Fig. 1C**). It is reasonable to believe that the occupation of MDR (including, but not limited to the co-resistance to tigecycline and polymyxin, two last-resort anti-infection options) by pMS8345A can be a serious risk in the clinic sector, once it successfully enters and further disseminates across pathogenic species.

### Genomic analyses of *tet*(X6)-containing MDR region

The linear genome alignment of MDR plasmids revealed that pMS8345A displays high level of similarity to at least three other resistance plasmids (**Fig. 1A**). Namely, they include i) the *mcr-1*-harboring plasmid pHNSHP45-2 with 99.75% identity and 87% query coverage (Acc. no.: KU341381), ii) and a tetracycline resistance plasmid of *Yersinia pseudotuberculosis*, pYps.F1 with 100% identity and 74% query coverage (Acc. no.: LT221036), and iii) a typical IncHI2-group, *mcr-1*-carrying plasmid pDJB-3 with 99.74% identity and 63% query coverage (Acc. no.: MK574666). Unlike that the *mcr-1*-lacking plasmid pYps.F1 exists in *Y. pseudotuberculosis*, all the other three *mcr-1*-harboring plasmids disseminate in different clones of *E. coli* with varied sequence types (**Fig. 1A**). In brief, i) pMS8345A is detected in ST95, a globally-distributed clone having the relevance to bacterial bloodstream infections and neonatal meningitis ^24^; ii) pHNSHP45-2 is recovered from an intensive pig farm ^29^; and iii) pDJB-3 is recently determined by our group to occur in ST165 in an pig farm, a rare sequence type (**Fig. 1A**).

Among them, the organization of MDR differs greatly (**Fig. 1**). Unlike the pMS8345A having both *tet*(X6)-positive MDR (**Fig. 1B**), and the *mcr-1*-containing cassette (**Fig. 1C**), the plasmid pDJB-3 has *mcr-1*, but not MDR region (**Fig. 1A**). In contrast, the plasmid pYps.F1 carries the MDR region, but not *mcr-1* (**Fig. 1A**). Genetic analysis elucidated that three class I integrons are located in the MDR region. Given that a pool of gene cassettes can be integrated, the majority of which encode resistance to antibiotics ^30^, Class I integron facilitates the global spread of AMRs ^31^. In particular, the MDR region of pMS8345A seems structurally unusual (**Fig. 1B**). First, it comprises a cluster of transposons and insert sequences (*IS*) with a boundary of two integrases (one is an integrase-encoding gene, *int1*, on the forward strand, and the other denotes a truncated version of *int1* on the reverse strand, **Fig. 1B**); Second, *int1* is adjacent to an integron-associated recombination site *attI*, and then followed by several *attC* sites (**Fig. 1B**); Third, the occurrence of Tn*2* and Tn*As1* (two copies of Tn*3* family transposons) in the pMS8345A MDR region implies an association with its mobility (**Fig. 1B**); Fourth, the multiple *IS* elements located within the MDR region (namely *bla*_CTX-M-1_ carried on an IS*Ecp1*, the operon of *strA*-*strB* adjacent to an IS*26*-IS*1133* structure ^32^, the *aac*(*3*)*-IIa/tmrB* plus *bla*_TEM-1B_ genes neighbored by IS*26*), facilitate the formation of transposons via the recombination events (**Fig. 1B**); Fifth, circular gene cassettes [such as *arr-2*/*ere*(*A*)/*aadA1*] are presumably integrated by site-specific recombination between *attI* and *attC*, a process mediated by the integron integrase (**Fig. 1B**). Along with class I integron (**Fig. 1B**), the fact that the GC content (37.8%) of *tet*(X6) is far less than the average GC% (46.29%) of the pMS8345A, allowed us to speculate that it is probably acquired via gene horizontal transfer. Sequence alignment reveals that the *tet*(X6)-positive MDR region in pMS8345A is highly similar to the MDR region in p15C38-2 (**Fig. 1B**). In brief, *tet*(X6) and *sul1* are downstream of a class 1 integron carrying the *aadA* family of resistance genes (*aadA22* in pMS8345A and *aadA12* in p15C38-2). p15C38-2 harbors a Tn*3*-like transposon, Tn*As2* with 85% identity.

### Functional insights into Tet(X6) tigecycline resistance

In addition to the pMS8345A plasmid, the *tet*(X6)-based *in silico* search uncovers two more *tet*(X6)-containing contigs (**Fig. S2**), namely TG_7F_Contig_3 (2575bp, Acc. no.: KU547185) and TE_7F_Contig_3 (2588bp, Acc. no.: KU547176). More intriguingly, the two contigs derive from uncultivated bacterium from latrine, in EI Salvador, 2012 (**Fig. 2**). It seems likely that *tet*(X6) appears earlier than that of *tet*(X4) initially detected in a contig of *K. pneumoniae* (4069bp, Acc. no.: NQBP01000050) from Thailand, 2015 ^20^. Further database mining suggests a number of *tet*(X) homologs [designated Tet(X7) to Tet(X13)] that are similar to *tet*(X6) at the identity ranging 91.27% to 97.62%. To relieve the confused nomenclature, we renamed the two redundant genes *tet*(X5) and *tet*(X6) appropriately (**Fig. 2**), which are chromosomally encoded in certain species like *Myroides* ^33^ and *Proteus* ^33-35^. Different from the pMS8345A plasmid-borne *tet*(X6), the designation of *tet*(X6) from four different species [*Proteus genomospecies* T60 ^34^, *P. cibarius* strain ZF2 ^35^, *Acinetobacter* (contig) ^33^ and *A. johnsonii* (contig) ^33^] is identical to that of *tet*(X12) we proposed here. Accordingly, the pAB17H194-1 plasmid-encoding *tet*(X5) in *Acinetobacter pittii* strain AB17H194 (Acc. no.: CP040912) is relabeled with *tet*(X14). The three inconsistent *tet*(X6) genes that arise separately from *P. cibarius* [contig, Acc. no.: WURM01000016] *P. mirabilis* [contig, Acc. no.: WURR01000048], and *M. phaeus* [genome, Acc. no.: CP047050] ^33^, were re-assigned with three distinct variants, namely *tet*(X15), *tet*(X16), and *tet*(X17). Phylogeny of these Tet(X) enzymes illustrates an ongoing Tet(X) family of resistance determinants (**Fig. 2**), raising a possible ancestor shared amongst these *tet*(X) variants.

**Figure 2.**
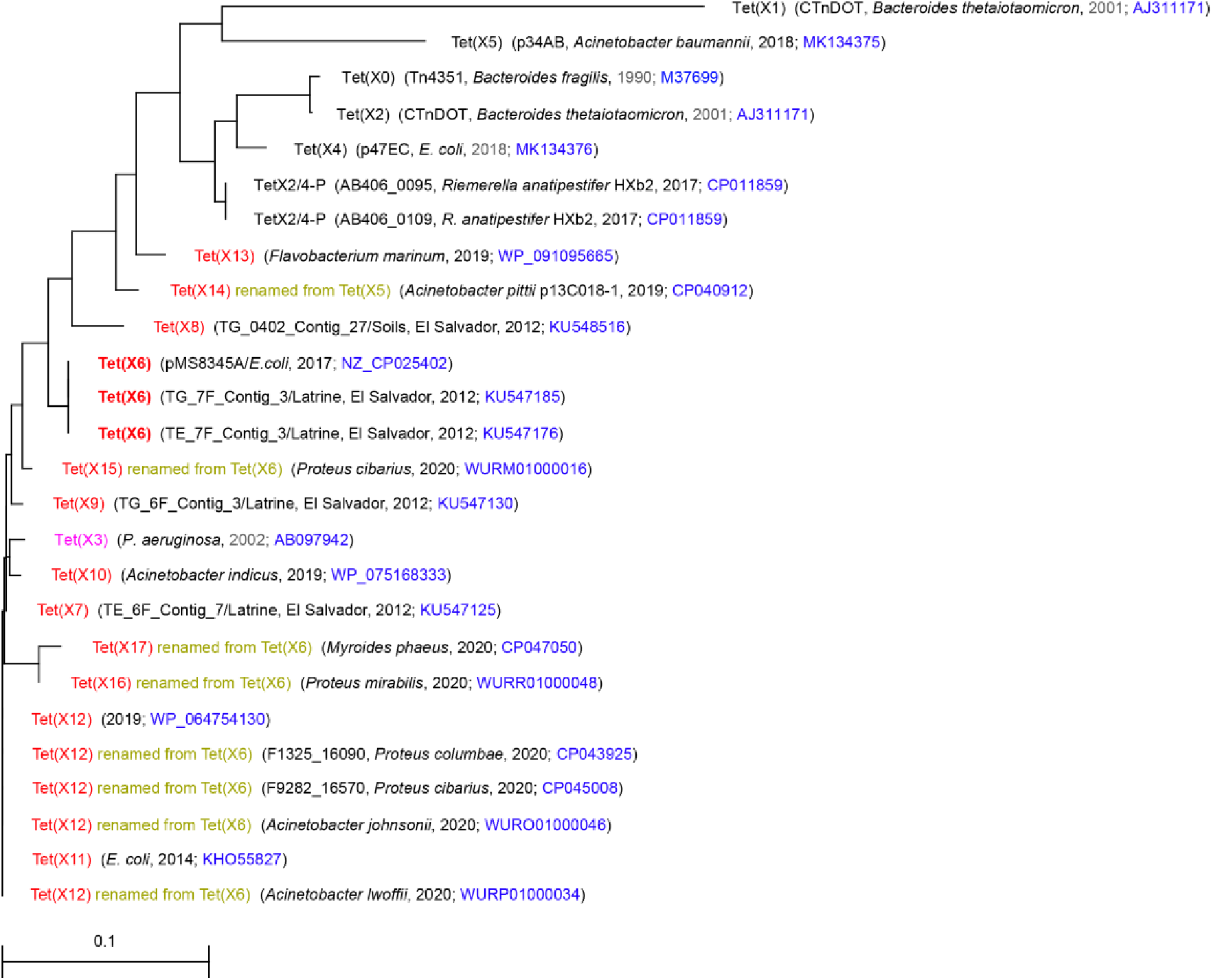
Phylogeny of Tet(X6) The protein sequences of Tet(X) variants were selected from NCBI database, and the accession numbers were labeled accordingly on the graph. Clustal Omega (https://www.ebi.ac.uk/Tools/msa/clustalo/) was applied to generate the phylogenetic tree, whose final output is built with TreeView (https://www.treeview.co.uk/). In addition to Tet(X3) (colored pink), all the twelve newly-renamed TetX variants (X6 to X17) are labeled in red. Besides the pMS8345A plasmid producing TetX6, two uncultured bacterium contigs (TE_7F_Contig_3 and TG_7F_Contig_3) from Latrine, EI Salvador, were found to carry an intact version of *tet*(X6). Of note, a number of *tet*(X) variants in duplicated and/or redundant nomenclature were corrected.

Given that i) the statement by He *et al*. ^18^ that TetX3 of *Pseudomonas* (Acc. no.: AB097942) is not active, is argued by our recent study ^20^; and ii) as a new variant, Tet(X6) displays 96.03% identity to the *Pseudomonas* Tet(X3) (**Fig. S3**), integrative evidences are highly demanded for the functional assignment of Tet(X6) in the context of tigecycline resistance. Therefore, we cloned *tet*(X6) into an arabinose-inducible pBAD24 expression vector and test its function in the strain MG1655 of *E. coli*. As predicted, the presence of *tet*(X6) can restore the growth of its recipient strain on LB agar plates with tigecycline (16 to 32μg/ml, **Fig. S4A**). This level of resistance is almost as same as *tet*(X3) does, but slightly lower than that of Tet(X4) (**Fig. S4A**). As predicted, structural modeling of Tet(X6) presents a substrate-loading channel (**Figs 3A-B**). Similar to the scenario with Tet(X4) ^20^, it consists of a tigecycline substrate-binding motif (**Figs 3C** and **E**) and a FAD cofactor-occupied cavity (**Figs 3D** and **F**). As for Tet(X6), the substrate-binding requires the cooperation of five residues E182, R203, H224, G226 and M365 (**Table 1** and **Fig. 3E**). Similarly, the FAD cofactor is surrounded with the following six residues E36, R37, R107, D301, P308 and Q312 in Tet(X6) (**Table 1** and **Fig. 3F**). Except that the substitution of H224T occurs in Tet(X1) [and/or H234Y in Tet(X5), in **Fig. S3**], all the aforementioned residues are relatively-conserved across the newly-proposed family of Tet(X) tigecycline-inactivating enzymes (**Table 1**). Among them, a number of residues have been functionally verified. In the case of Tet(X4), two of 5 substrate-binding cavity (H231 and M372) and three of 6 FAD-interactive residues (E43, R114, and D308) somewhat play roles in the phenotypic tigecycline resistance ^20^. Structure-guided alanine substitution of Tet(X6) revealed that i) two of 5 tigecycline-binding residues (H224A and M365A) partially determine its phenotypic resistance to tigecycline (**Fig. 3G**); and ii) three of 6 FAD-surrounding residues (E36A, R107A, and D301A) give differential level of impact on its resultant tigecycline resistance (**Fig. 3H**). Thus, this result represents a functional proof for Tet(X6) as a new member of the expanding family of Tet(X) enzymes that have a role in the action of tigecycline degradation.

**Table 1.** Substrate-loading cavities across Tet(X)-type enzymes The substituted residues are indicated in yellow background. *Tet(X14) is renamed from Tet(X5) of *Acinetobacter pittii* p13C018-1 [acc. no.: CP040912]; Tet(X15) is relabeled from Tet(X6) of *Proteus cibarius* [acc. no.: WURM01000016]; Tet(X16) replace the redundant Tet(X6) from *Proteus mirabilis* [acc. no.: WURR01000048]; and Tet(X17) is corrected from Tet(X6) of *Myroides phaeus* [acc. no.: CP047050].

| Enzymes/Length | Substrate-loading cavities |  |  |  |  |  |  |  |  |  |  |
| --- | --- | --- | --- | --- | --- | --- | --- | --- | --- | --- | --- |
|  | FAD-binding sites |  |  |  |  |  | Tetracycline-recognizable sites |  |  |  |  |
| Tet(X0), 388aa | E46 | R47 | R117 | D311 | P318 | Q322 | Q192 | R213 | H234 | G236 | M375 |
| Tet(X1), 379aa | E36 | R37 | R107 | D302 | P309 | Q313 | Q182 | R203 | T224 | G226 | M366 |
| Tet(X2), 388aa | E46 | R47 | R117 | D311 | P318 | Q322 | Q192 | R213 | H234 | G236 | M375 |
| Tet(X3), 378aa | E36 | R37 | R107 | D301 | P308 | Q312 | Q182 | R203 | H224 | G226 | M365 |
| Tet(X4), 385aa | E43 | R44 | R114 | D308 | P315 | Q319 | Q189 | R210 | H231 | G233 | M372 |
| Tet(X5), 386aa | E46 | R47 | R117 | D310 | P317 | Q321 | Q192 | R213 | Y234 | G236 | M373 |
| Tet(X6), 378aa | E36 | R37 | R107 | D301 | P308 | Q312 | Q182 | R203 | H224 | G226 | M365 |
| Tet(X7), 378aa | E36 | R37 | R107 | D301 | P308 | Q312 | Q182 | R203 | H224 | G226 | M365 |
| Tet(X8), 378aa | E36 | R37 | R107 | D301 | P308 | Q312 | Q182 | R203 | H224 | G226 | M365 |
| Tet(X9), 378aa | E36 | R37 | R107 | D301 | P308 | Q312 | Q182 | R203 | H224 | G226 | M365 |
| Tet(X10), 387aa | E45 | R46 | R116 | D310 | P317 | Q321 | Q191 | R212 | H233 | G235 | M374 |
| Tet(X11), 378aa | E36 | R37 | R107 | D301 | P308 | Q312 | Q182 | R203 | H224 | G226 | M365 |
| Tet(X12), 387aa | E45 | R46 | R116 | D310 | P317 | Q321 | Q191 | R212 | H233 | G235 | M374 |
| Tet(X13), 388aa | E46 | R47 | R117 | D310 | P317 | Q321 | Q192 | R213 | H234 | G236 | M373 |
| Tet(X14) *, 388aa | E46 | R47 | R117 | D311 | P318 | Q322 | Q192 | R213 | H234 | G236 | M375 |
| Tet(X15) *, 387aa | E45 | R46 | R116 | D310 | P317 | Q321 | Q191 | R212 | H233 | G235 | M374 |
| Tet(X16) *, 387aa | E45 | R46 | R116 | D310 | P317 | Q321 | Q191 | R212 | H233 | G235 | M374 |
| Tet(X17) *, 387aa | E45 | R46 | R116 | D310 | P317 | Q321 | Q191 | R212 | H233 | G235 | M374 |
| Tet(X2/4)-P, 388aa | E46 | R47 | R117 | D310 | P317 | Q321 | Q192 | R213 | H234 | G236 | M373 |

*Myroides phaeus* [acc. no.: CP047050].

**Figure 3.**
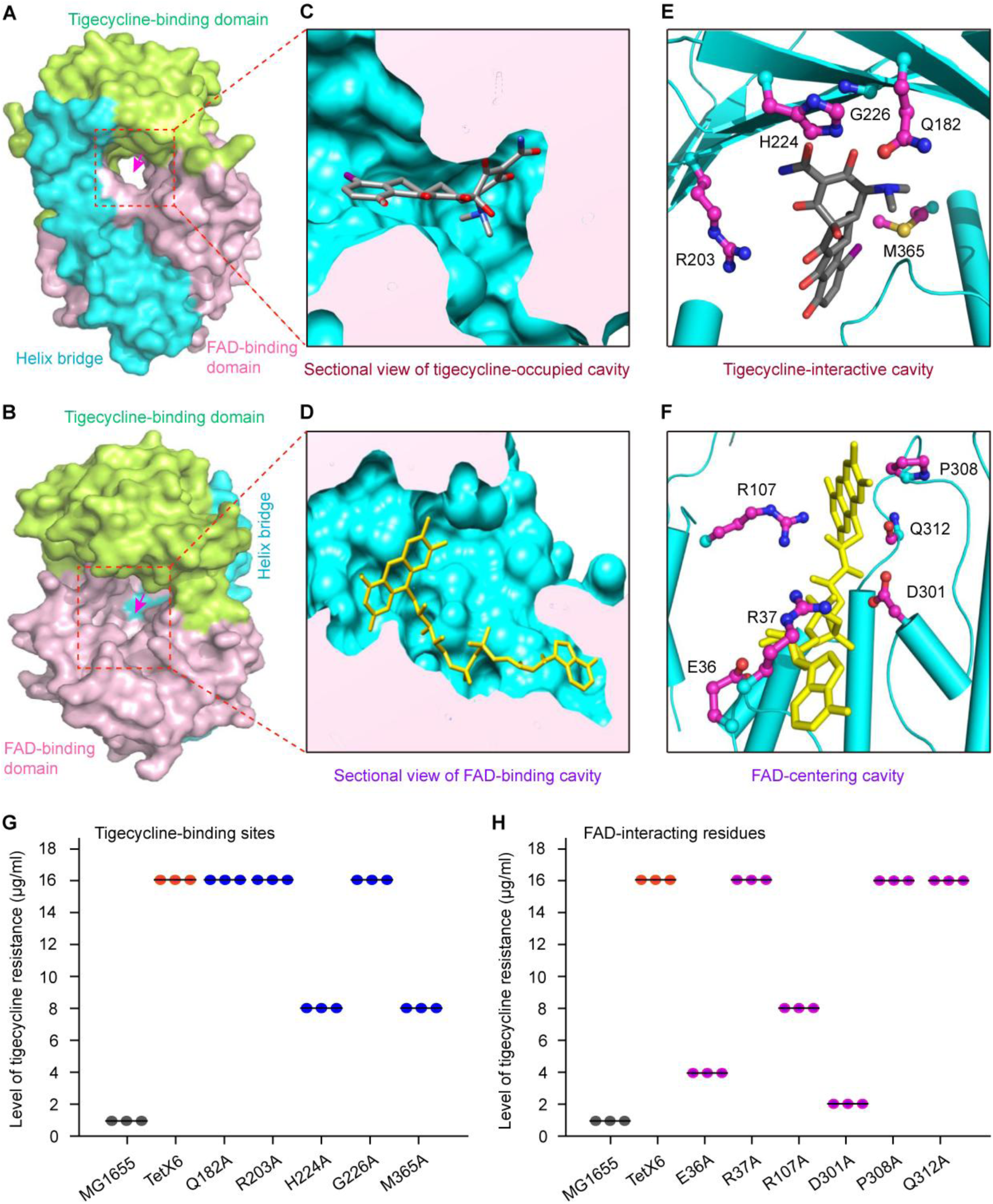
Structural and functional analysis of the substrate tigecycline-loading tunnel in Tet(X6) Surface structure of Tet(X6) with tigecycline-loading tunnel in the front view (**A**) and the rotated view (**B**). The substrate-loading tunnel in open state is indicated with an arrow. **C.** Sectional view of tigecycline-occupied cavity in Tet(X6) protein **D.** Sectional view of FAD-binding cavity in Tet(X6) enzyme **E.** An enlarged view of tigecycline-binding cavity in Tet(X6) **F.** Structural snapshot of the FAD cofactor-binding motif in Tet(X6) The critical residues are labeled. The molecule of tigecycline is colored in cherry red, and the FAD molecule is labeled in purple. **G.** Use of site-directed mutagenesis to assay the five tigecycline-binding residues in Tet(X6) **H.** Structure-guided alanine substitution analyses of FAD-interacting residues in Tet(X6) Negative control is the *E. coli* MG1655 alone, and positive control refers to the MG1655 expressing the wild-type of *tet*(X6) (**Table S1**). The pBAD24-driven expression of Tet(X6) and its derivatives is triggered by the addition of 0.2% arabinose. Three independent assays were conducted, each of which is indicated with a dot.

### Action of inactivation of tigecycline by Tet(X6)

To further elucidate biochemical mechanism of Tet(X6)-catalyzed tigecycline destruction (**Fig. 4**), we integrated an *in vivo* approach of microbial bioassay (**Fig. 4A**) with the *in vitro* system of enzymatic reaction (**Figs 4B-D**). This tigecycline bioassay we developed is dependent on the indicator strain DH5α of *E. coli* in that it was verified to be tigecycline susceptibility (**Fig. 4A**). As predicted, a zone of bacterial inhibition was clearly seen to surround a paper disk of the blank control, on which 2.5µg/ml tigecycline is spotted (**Fig. 4A**). A similar scenario was also seen with the negative control, i.e., the supernatant from *E. coli* MG1655 having the empty vector pBAD24 alone (**Fig. 4A** and **Table S1**). In contrast, the zone of tigecycline inhibition disappears around the paper disk containing supernatants of *E. coli* MG1655 expressing either *tet*(X6) or its homologous gene *tet*(X3) (**Fig. 4A**). This highlighted an *in vivo* role of *tet*(X6) [and/or *tet*(X3)] in the degradation of tigecycline.

**Figure 4.**
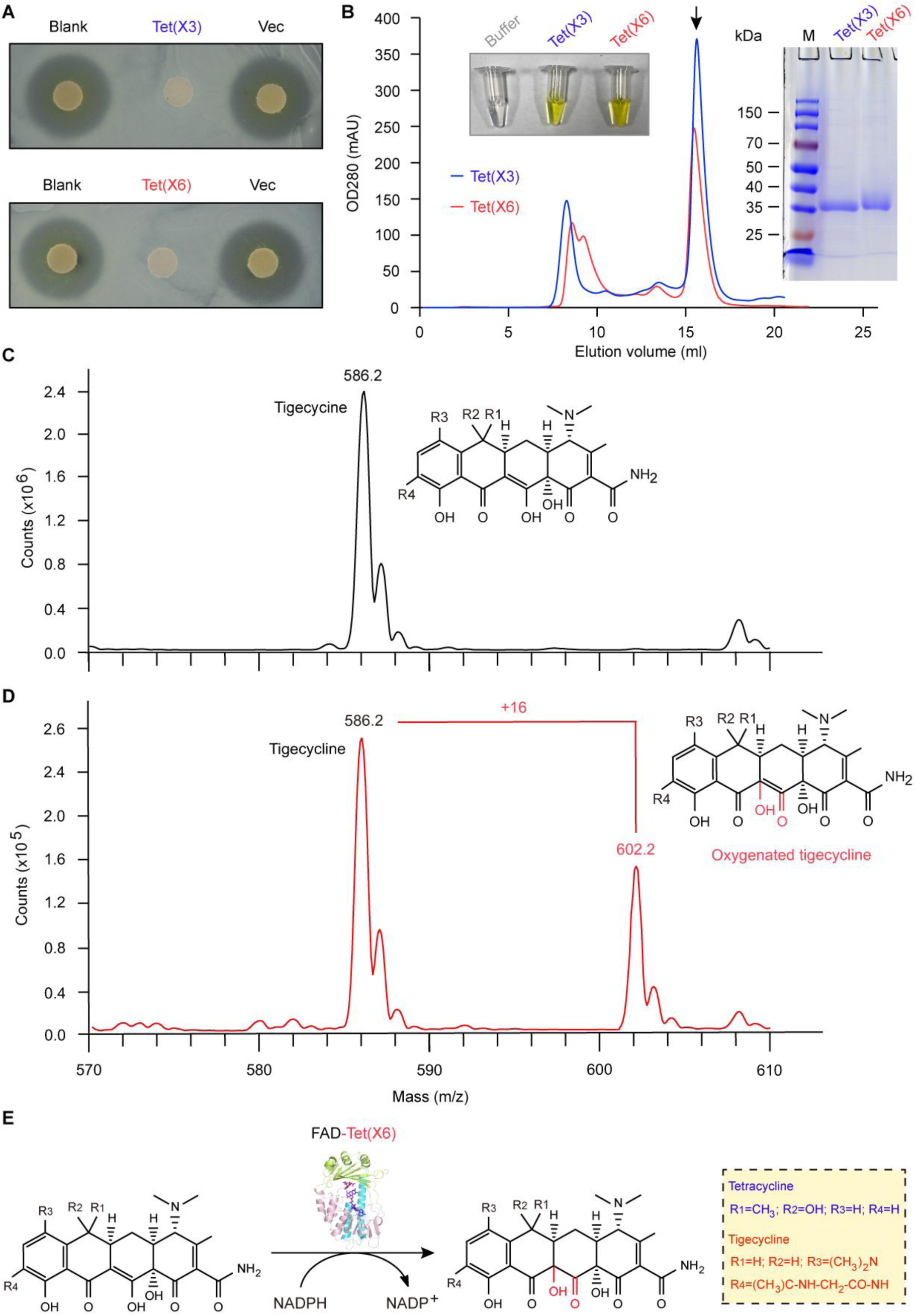
Action and mechanism for Tet(X6) tigecycline resistance. **A.** The bioassay for Tet(X6) [and Tet(X3)]-mediated destruction of tigecycline The presence of tigecycline on paper disks leads to the appearance of a transparent zone of microbial inhibition. In contrast, the inhibition zone is absent upon the destruction of tigecycline by Tet(X6) [and/or Tet(X3)]. **B.** Gel filtration analysis of the purified Tet(X6) resistance enzyme To address the solution structure of Tet(X6) and Tet(X3), the method of size exclusion chromatography was applied. The protein samples were loaded into Superdex 200/300GL size exclusion column (GE Healthcare, USA). The eluted protein of interest (indicated with an arrow) is visualized in the right-hand inside gel (15% SDS-PAGE). The almost-identical elution volume (∼16ml) of Tet(X6) and Tet(X3) is generally consistent with the apparent size of molecular weight ∼36kDa in the inside PAGE gel, validating the monomeric form. Of note, the yellow color of Tet(X6) [Tet(X3)] solution is given in inside gel on the left hand, indicating the presence of a FAD cofactor-bound protein form. LC/MS identification of tigecycline (**C**) and its oxygenated product by Tet(X6) enzyme (**D**) Compared with the peak of tigecycline at 586.2 (m/z), the oxygenated derivative of tigecycline is reflected by a unique peak of 602.2 (m/z) in that it is added with an oxygen atom. **E.** A chemical reaction model that Tet(X6) destructs/inactivates tigecycline It was adapted appropriately from Xu *et al*. ^20^ with permission. FAD-Tet(X6) denotes Tet(X6) enzyme in the form of binding FAD cofactor. Chemical structures were given with ChemDraw. Designations: blank, the LBA media with only tigecycline; vec, the vector of pET21a; LC/MS, liquid chromatography mass spectrometry; FAD, flavin adenine dinucleotide; NADP^+^, the oxidized form of nicotinamide adenine dinucleotide phosphate; NADPH, the reduced form of nicotinamide adenine dinucleotide phosphate.

Subsequently, we produced the recombinant forms of Tet(X6) and its homolog Tet(X3) and examined their enzymatic activities *in vitro*. Notably, Tet(X6) and Tet(X3) protein consistently gives yellow in solution (**Figs 4B** and **5B**), hinting a possibility of being occupied with a FAD cofactor (**Fig. 5A**). Indeed, optical absorbance spectroscopy demonstrated the presence of Tet(X6)-bound FAD cofactor (**Figs 5A-B**). Gel filtration analysis indicated that both Tet(X6) and Tet(X3) display the solution structure of being a monomer (**Fig. 4B**). This generally agrees with the apparent molecular mass (∼36kDa) seen in the SDS-PAGE (**Fig. 4B**). The identification of polypeptide fingerprint with liquid chromatography (LC)/mass spectrometry allowed us to further study the catalytic action of Tet(X6) [Tet(X3)] using the *in vitro* reconstituted system of tigecycline oxygenation (**Figs S5A-B**). As expected, LC/MS-based detection of the substrate tigecycline showed a unique peak at the position of 586.2 m/z (**Fig. 4C**). In particular, the reaction mixture of Tet(X6) [Tet(X3)] consistently gave two distinct peaks in the spectrum of LC/MS (**Figs 4D** and **S6**). Namely, they correspond to a peak of substrate tigecycline (586.2 m/z), and an additional peak assigned to its oxygenated product of tigecycline at the position of 602.2 m/z (**Figs 4D** and **S6**). Notably, the method of double-reciprocal plot (**Figs 5C-D**) was exploited to measure the kinetic parameters (**Fig. 5E**) of Tet(X6) enzyme for the reactant tigecycline. As a result, Km of Tet(X6) was calculated to be 42.6±4.3 (**Figs 5E-F**), which is comparable to those of Tet(X2), Tet(X4) and Tet(X5) (**Fig. 5F**). This finding is consistent with an observation with the newly-identified Tet(X4) by He and coauthors ^18^.

**Figure 5.**
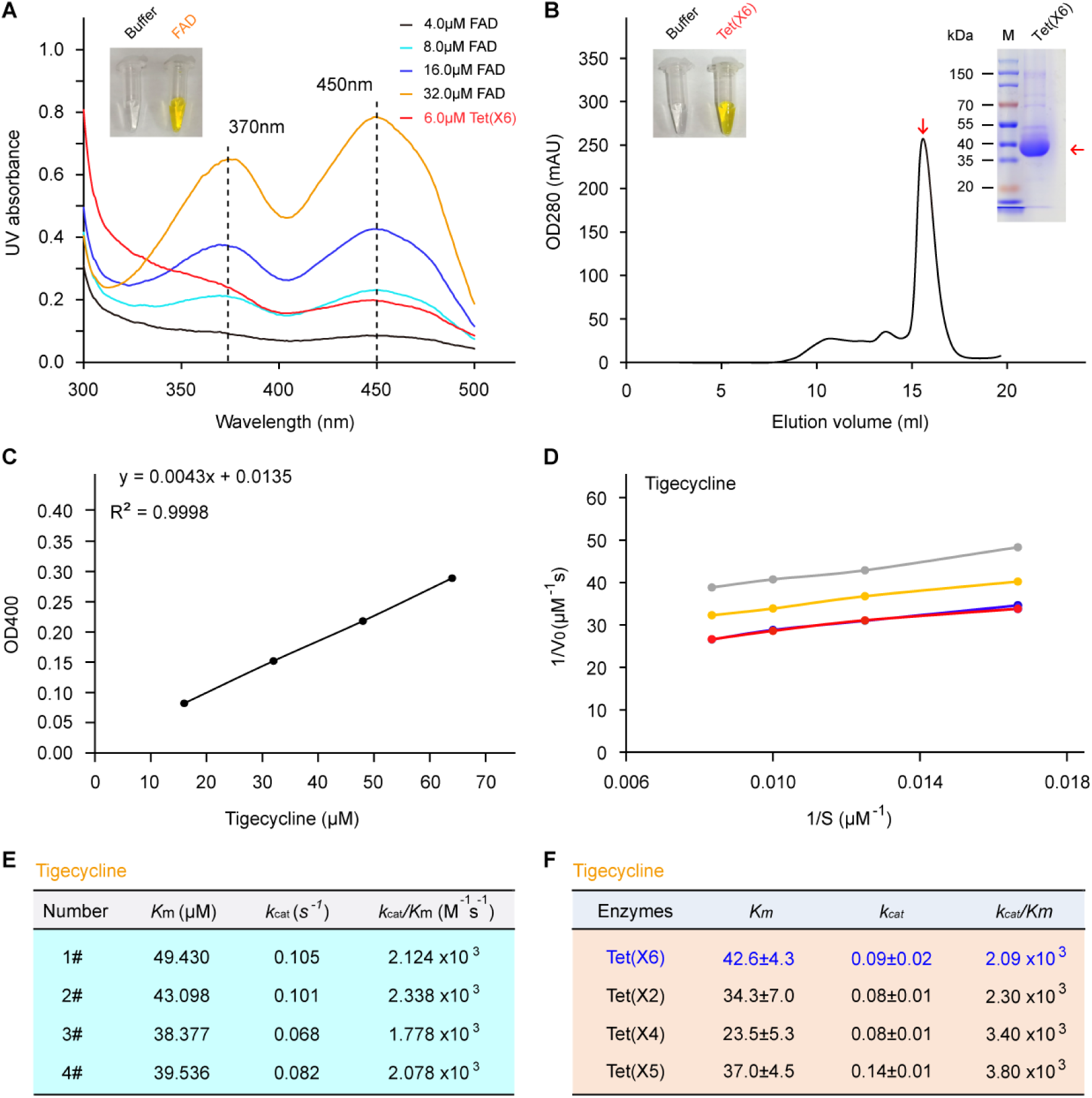
Kinetic characteristics of Tet(X6) enzyme. **A.** Use of optical absorbance spectroscopy to detect the Tet(X6)-bound FAD cofactor The FAD solution (positive control) with yellow color (inside gel), features with two unique peaks at the wave-lengths of 370nm and 450nm. A similar scenario was also seen with the sample of Tet(X6) protein. **B.** Purity of the recombinant Tet(X6) protein judged with gel filtration The purified protein of Tet(X6) with the yellow color (inside gel on left hand), was separated with 15% SDS-PAGE (inside gel on right hand). The gel filtration was developed with the Superdex 200 column run on AKTA Pure. **C.** The standard curve of absorption of tigecycline at the wavelength of 400nm. The slope of the line is described the symbol “ε”. As for tigecycline, ε_400_=4300 M^-1^ cm^-1^. Use of double-reciprocal plot (**D**) to measure the kinetic parameters (**E**) of Tet(X6) enzyme Four independent experiments were conducted to generate the above double-reciprocal plots, giving kinetic parameters. **F**. Kinetic constant of Tet(X6) is comparable with the counterpart of the other three known Tet(X) enzymes, namely Tet(X2), Tet(X4) and Tet(X5) The value of Tet(X6) arising from the data (**panel D**) are expressed in averages±SD. All the values of other three enzymes were reported by He and coworkers ^18^. Designations: K_m_, Michaelis-Menten constant; V_max_, the maximum velocity of enzymatic reaction; K_cat_, catalytic constant; V_0_, the initial velocity; SD, standard deviation.

As Forsberg *et al*. ^17^ stated, similar scenarios were also seen with both Tet(X3) and Tet(X6) (**Fig. S7**), which is evidenced by the fact that liquid culture of *tet*(X3) [and/or *tet*(X6)]-bearing *E. coli* gives dark (**Fig. S7**). Unlike that the negative-control strain MG1655 with empty vector pBAD24 alone displays a big inhibition circle, E-test of tigecycline showed that expression of *tet*(X3) [or *tet*(X6)] renders the recipient strains significantly antagonistic to the tigecycline challenge (**Fig. S8**). Thereafter, we formulated a working model that Tet(X6) exploits a FAD cofactor to oxygenate/destruct the last-line antibiotic tigecycline (**Fig. 4E**). It seems likely that this chemical reaction proceeds via a ‘ping-pong’ action. However, this hypothesis requires further experimental evidence.

### Variation in the *mcr-1*-containing cassettes

Sequence analysis of *mcr-1*-bearing elements from the three IncHI2-type plasmids (pMS8345A, pSA186_MCR1, and pDJB-3) reveals the core structure of “IS*Apl1-mcr-1*” (**Fig. 1C**). Unlike the plasmid of pDJB-3 containing a cassette of “IS*Apl1*-*mcr-1-pap2-*IS*Apl1*”, the pSA186_MCR1 plasmid possesses an inactivated *pap2* inserted with an inverted copy of IS*Apl1* (**Fig. 1C**). As a member of the IS*30* family, the IS*Apl1* can transpose into its target by formatting a synaptic complex between an inverted repeat (IR) in the transposon circle and an IR-like sequence in the target ^36^. After the initial formation of this composite transposon, one or both copies of IS*Apl1* might be lost. As such, this loss might improve the stability of *mcr-1* in a diverse range of plasmids and then intensify its spread of *mcr-1* ^37^. Therefore, we favor to believe this model that pMS8345A having “IS*Apl1-mcr-1*” alone (**Fig. 1C**) proceeds the loss of its downstream IS*Apl1* following the transposition. Not surprisingly, the *E. coli* strain carrying *mcr-1* gives the minimum inhibitory concentration (MIC) at 4.0μg/ml. In addition, functional expression of a single *mcr-1* allows the polymyxin-susceptible recipient strain of *E. coli* MG1655 to appear on the LB agar plate with colistin of up to 16μg/ml (**Fig. S4B**). Evidently, these data demonstrated that both *tet*(X6) and *mcr-1* are actively co-carried by a single plasmid in an epidemic ST95 clone of pathogenic *E. coli*.

### Physiological alteration by Tet(X6) and MCR-1

To address physiological consequence of Tet(X6) and MCR-1, we separately examined the pool of intracellular reactive oxygen species (ROS) and various growth curve-based metabolic fitness in an array of different *E. coli* strains (**Figs 6** and **7A**). As illustrated in the assay of fluorescence activated cell sorting (FACS), the cytosolic ROS level in the M1655 with empty vector alone was relatively low (**Fig. 6A**). Similar scenarios were also seen in derivatives of the MG1655 strain, regardless of the presence of *mcr-1* (**Fig. 6D**), *tet*(X6) (**Fig. 6E**), and even both (**Fig. 6F**). As a consequence, the level of intracellular ROS was increased greatly upon its exposure to either colistin (**Figs 6B** and **J**) or tigecycline (**Figs 6C** and **J**). The expression of *mcr-1* effectively prevented the colistin-stimulated ROS formation (**Figs 6G** and **J**). Similarly, the presence of *tet*(X6) robustly interfered the ROS production triggered by tigecycline (**Figs 6H** and **J**). In fact, the addition of both colistin and tigecycline only gave slight increment of ROS accumulation in the MG1655 strain co-harboring *mcr-1* and *tet*(X6) (**Figs 6I-K**). Therefore, Tet(X6) attenuates the tigecycline-induced ROS generation as does MCR-1 in response to colistin (**Fig. 6**).

**Figure 6.**
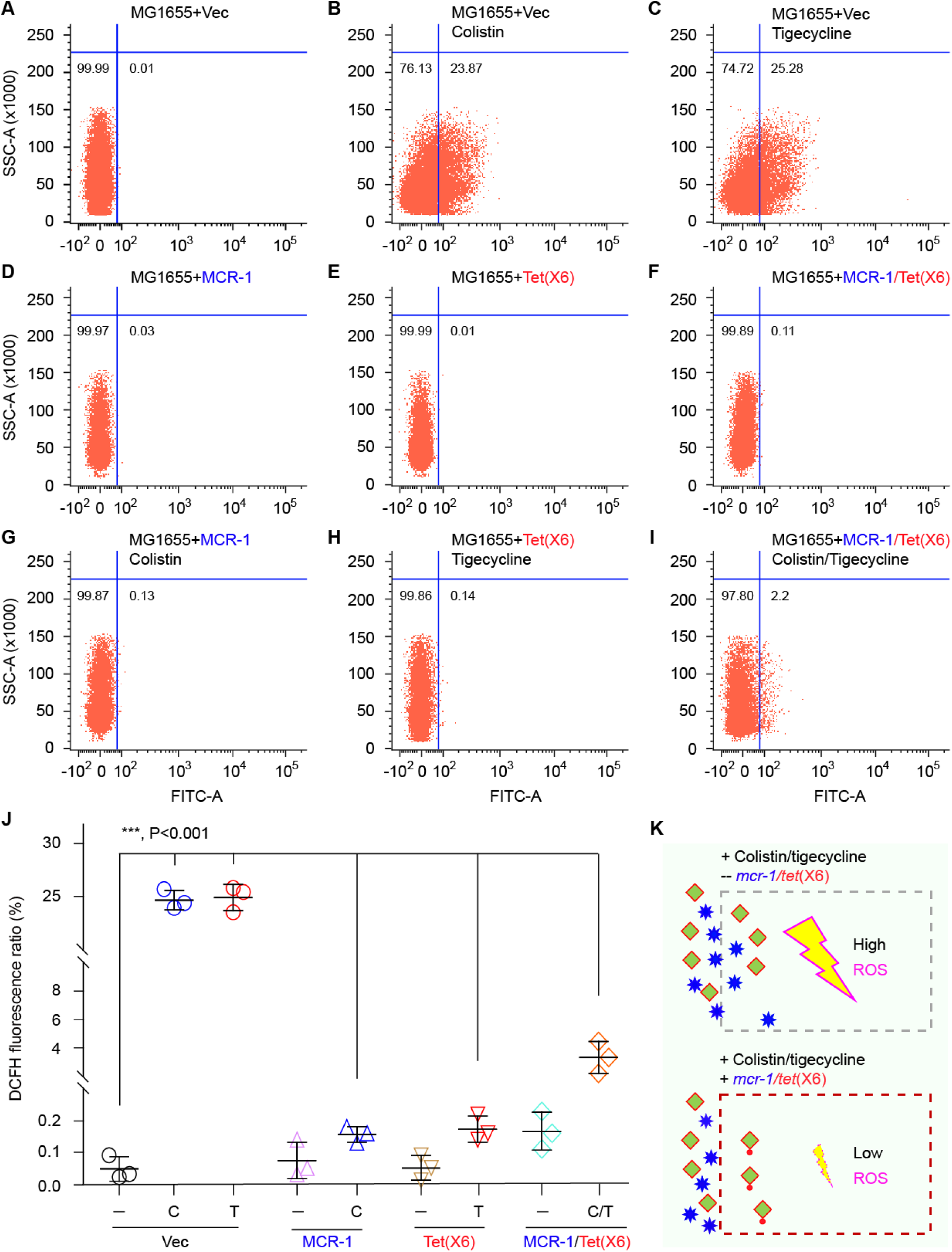
Use of flow cytometry to measure intracellular ROS level induced by colistin and/or tigecycline. **A.** FACS analysis of basal level of ROS in the negative strain *E. coli* MG1655 with empty vector alone Intracellular ROS level is boosted upon the addition of either colistin (**B**) or tigecycline (**C**) into the MG1655 strain Cytosolic ROS level of the MG1655 strains expressing *mcr-1* (**D**) or *tet*(X6) (**E**) **F.** Determination of ROS level in the MG1655 carrying both *mcr-1* and *tet*(X6) **G.** Colistin cannot stimulate the ROS production in the MG1655 strain expressing *mcr-1* **H.** Tigecycline cannot activates the ROS production in the MG1655 strain expressing *tet*(X6) **I.** The mixture of tigecycline and colistin cannot significantly alter the cytosolic ROS accumulation in the MG1655 strain co-harboring *mcr-1* and *tet*(X6) **J.** Flow cytometry-based determination of relative level of intracellular ROS in different *E. coli* MG1655 strains in response to colistin and/or tigecycline Along with Tukey-Kramer multiple comparisons post-test, the data is given using one-way analysis of variance (ANOVA). *p-value is less than 0.001. **K.** A scheme for colistin/tigecycline-induced accumulation of ROS, and its interfered formation by MCR-1/Tet(X6) Blue asterisk denotes colistin, and green square refers to tigecycline. The oxygenated form of tigecycline is indicated with green square attached with a red dot. The reactive oxygen species (ROS) are illustrated with the symbols of lightning. As for the induction of ROS, the two antibiotics used here denoted 0.2mg/ml colistin and 0.2mg/ml tigecycline. Abbreviations: FACS, fluorescence activated cell sorting; ROS, reactivated oxygen species.

As expected, the presence of plasmid-borne *mcr-1* can cause the delayed growth of its recipient host *E. coli* MG1655, whereas the empty vector not (**Figs 7B&D**). In agreement with earlier observations ^38-42^, this underscored that MCR-1 exerts significantly fitness cost in *E. coli*. In contrast, the expression of *tet*(X6) fails to trigger any detectable retardation of bacterial growth (**Fig. 7C**), indicating that Tet(X6)-causing metabolic burden/disorder is minimal. To further probe whether or not the crosstalk between Tet(X6) and MCR-1 occurs in *E. coli*, we engineered an *E. coli* strain that coharbors derivatives of two compatible plasmids [one arises from a low copy number, lactose promoter-driven pWSK129 ^43^, and the other is constructed from an arabinose-inducible pBAD24 with ampicillin resistance ^44^]. In fact, the two resistance enzymes MCR-1 and Tet(X6) are produced by lactose-inducible pWSK129::*mcr-1*, and arabinose-activating pBAD24::*tet*(X6), respectively (**Table S1**). Evidently, the coexistence of *tet*(X6) and *mcr-1* cannot exert any synergism on bacterial retarded growth (**Fig. 7E**).

**Figure 7.**
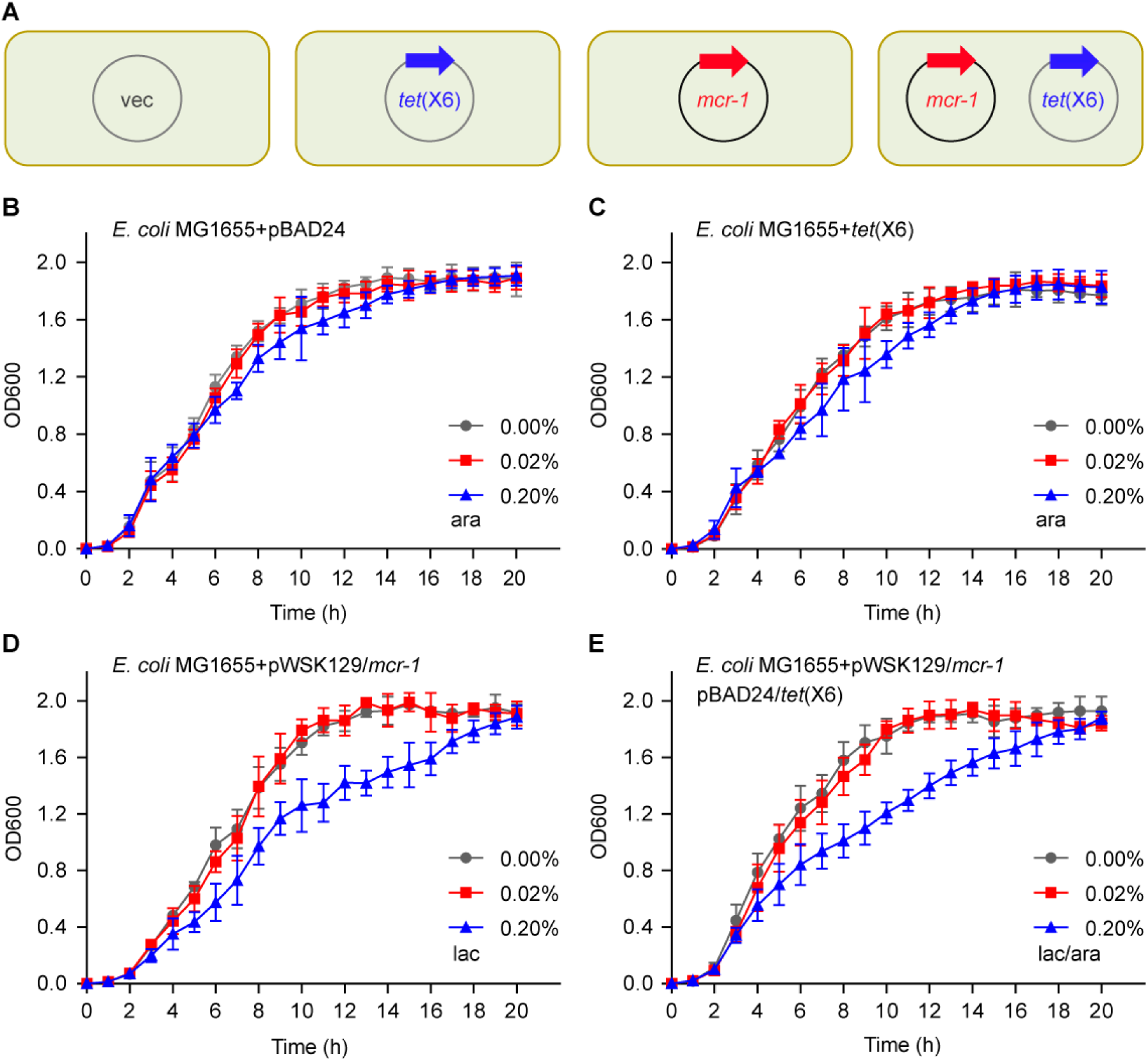
Use of growth curves to monitor metabolic fitness caused by *mcr-1* and *tet*(X6) in *E. coli*. **A.** Schematic representative of various *E. coli* strains carrying plasmid-borne *tet*(X6), *mcr-1* or both **B.** The empty vector exerts no effect on bacterial growth of the *E. coli* MG1655 **C.** Expression of *tet*(X6) causes slightly the delayed growth of the *E. coli* MG1655 **D.** Expression of *mcr-1* gives significantly metabolic fitness in its recipient host *E. coli* MG1655 slightly the delayed growth of the *E. coli* MG1655 **E.** Co-expression of *mcr-1* and *tet*(X6) cannot lead to synergistic fitness cost of the recipient strain The data was given in means ± SD from three independent plotting of growth curves. vec, pBAD24.

As recently performed with MCR-3/4, we also adopted an approach of LIVE/DEAD cell staining to analyze an array of engineered strains (**Fig. 8A**). Unlike that the negative control, MG1655 strains with vector alone are almost fulfilled with alive cells (**Figs 8B-C**), the *mcr-1*-producing strains contained around 30% dead cells (**Figs 8D-E**, and **J**). Consistent with that no retarded growth is associated with Tet(X6) (**Fig. 6C**), confocal microscopy assays illustrated that relatively-low level of DAED/LIVE ratio is present in the *tet*(X6)-carrying MG1655 (**Figs 8F-G**, and **J**). Not surprisingly, the co-expression of *mcr-1* and *tet*(X6) cannot promotes significant increment in the ratio of DEAD/LIVE cells (**Figs 8H-J**), when compared with the *mcr-1*-positive strains (**Figs 8D-E**). The remaining question to ask is whether or not Tet(X6) tigecycline resistance can crosstalk with MCR-1 colistin resistance in a given strain (**Fig. 9**). Thus, we designed such an *E. coli* strain FYJ4022 (**Table S1**), which co-harbors pWSK129::*mcr-1* and pBAD24::*tet*(X6). In this engineered strain, the expression of *mcr-1* is turned on by the addition of lactose, and *tet*(X6) is finely tuned by the supplementation of arabinose (**Fig. 9A**). The colistin resistance by MCR-1 was found to be independently of the presence of Tet(X6) (**Fig. 9B**), and vice versa (**Fig. 9C**). Further, MALDI-TOF mass spectrometry confirmed that the insusceptibility to polymyxin, arises from the PEA addition to lipid A by MCR-1, regardless of the presence of Tet(X6) in *E. coli* (**Figs 9D-E**).

**Figure 8.**
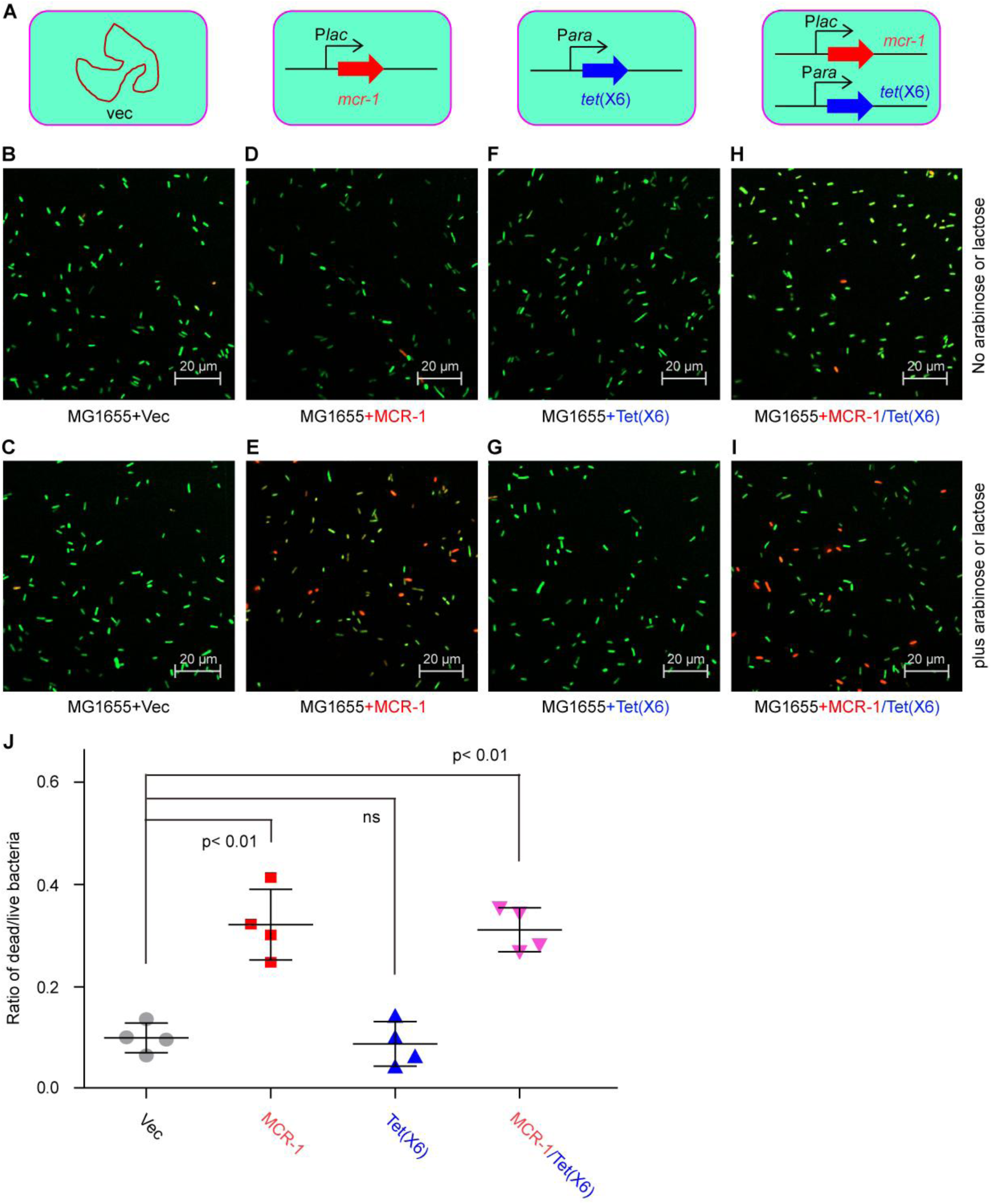
Confocal microscopy-based visualization for bacterial fitness cost caused by functional expression of *tet*(X6) (and/or *mcr-1*) in *E. coli*. **A.** Scheme for engineered strains of *E. coli* with inducible expression of *tet*(X6), *mcr-1*, or both **B&C** Bacterial viability of the negative control strains, *E. coli* MG1655 with the empty vector alone is not affected by the presence of the inducer arabinose and/or lactose **D&E** The ratio of DEAD/LIVE cells suggests that arabinose-induced expression of *mcr-1* gives appreciable level of fitness cost in *E. coli* **F&G** Regardless of the inducer lactose added, the MG1655 strains harboring *tetX6* is indistinguishable, when compared with those of negative control strains **H.** The *mcr-1* and *tet*(X6)-coharboring MG1655 does not display the phenotypic fitness cost, without the addition of neither arabinose nor lactose **I.** The co-expression of MCR-1 and Tet(X6) cannot exert synergistic effect of metabolic fitness **J.** Bacterial viability-based measurement of bacterial fitness costs in various MG1655 strains bearing either *mcr-1* or *tet*(X6) (or both) Bacterial viability was determined through cell staining with LIVE/DEAD kit, which is followed by imaging with confocal laser scanning microscopy. The color of green and red separately denotes alive and dead cells. Data was collected from four independent trials and evaluated using one-way analysis of variance (ANOVA) followed by Tukey–Kramer multiple comparisons post hoc test. **p-value is less than 0.01. ns, no significance.

**Figure 9.**
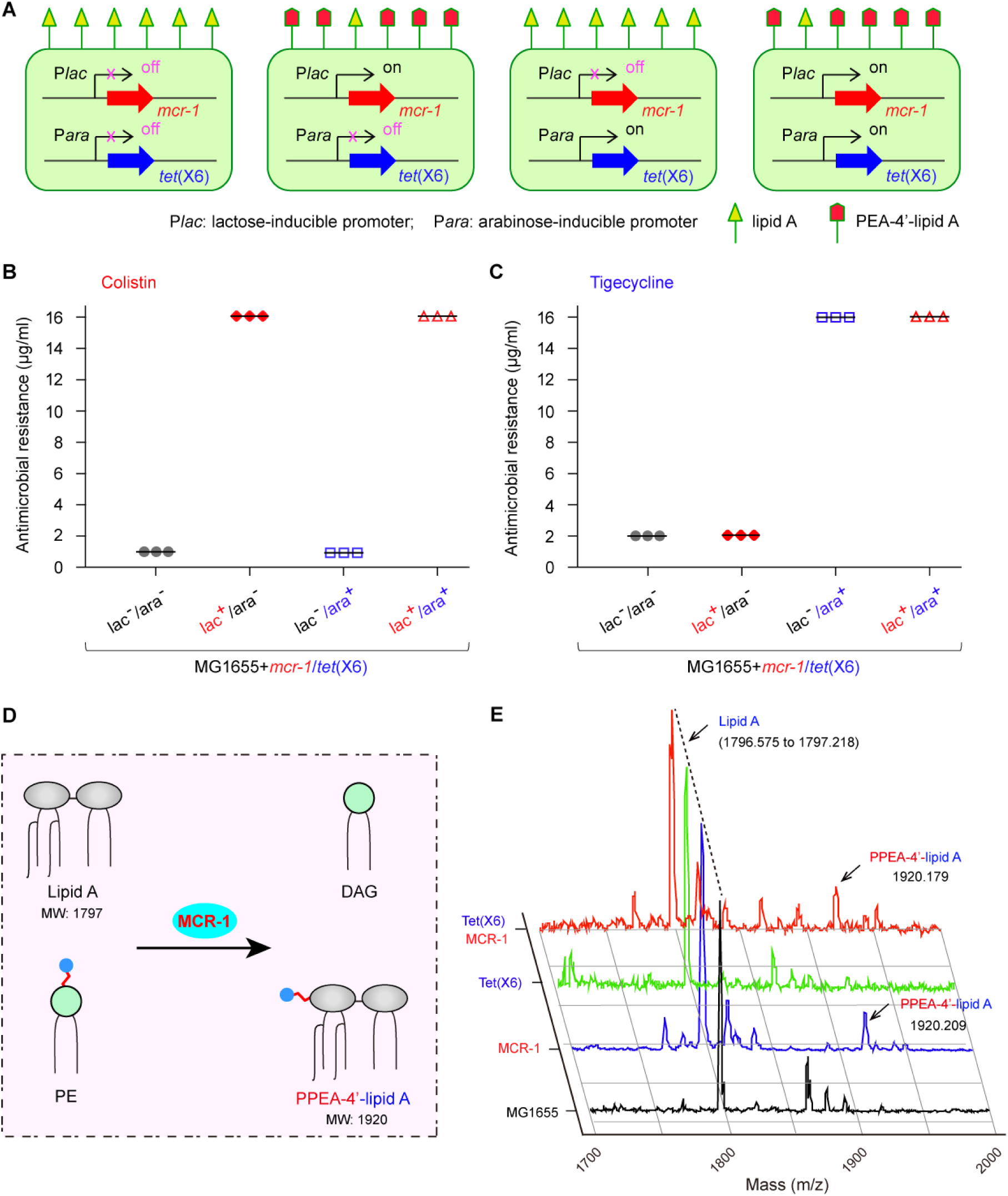
No crosstalk between MCR-1 and Tet(X6) in a given *E. coli* strain. **A.** Schematic representative of an engineered *E. coli* strain co-carrying *mcr-1* and *tet*(X6) in four different modes of expression A symbol of triangle denotes intact LPS-lipid A, whereas the symbol “” refers to the PPEA-4’-lipid A, a derivative of lipid A with the addition of PEA moiety. **B.** MCR-1 colistin resistance occurs independently of Tet(X6) tigecycline resistance **C.** Tet(X6) tigecycline resistance proceeds independently of MCR-1 colistin resistance **D.** Scheme for remodeling of bacterial lipid A by MCR-1 **E.** Structural identification of lipid A species from different strains of E. coli expressing a single *tet*(X6)/*mcr-1* or both The strain tested here denotes FYJ4022 (**Table S1**), which is MG1655 co-harboring pWSK129::*mcr-1* and pBAD24::*tet*(X6). The assays of susceptibility to colistin and tigecycline were performed with LB Agar plates with varied levels of antibiotics. The symbol “-” refers to no addition of arabinose or lactose, whereas “+” denotes addition of arabinose and/or lactose. Abbreviations: PE, Phosphatidylethanolamine; DAG, Diacylglycerol; PPEA, Phosphoethanolamine; MW, molecular weight.

Together, these data suggested that no synergism is detected in fitness costs caused by the lipid A modifier MCR-1 and the tigecycline-inactivating enzyme Tet(X6). Unlike that MCR-1 modifies bacterial lipid A, the initial target of the cationic antimicrobial peptide colistin receptor ^5^, Tet(X6) hydrolyzes the family of tetracycline and its derivatives, like tigecycline (**Fig. 4**) ^15,20^. The former action results in bacterial surface remodeling by MCR-1 through an addition of PEA moiety to 1(4’)-phosphate position of lipid A ^8,9^. Consequently, this might in part shape metabolic flux of the recipient microbe to balance *mcr-1* expression and bacterial survival stressed with colistin, producing the phenotypic fitness cost ^38^. In contrast, it seems likely that the destruction of tigecycline by the flavin-dependent Tet(X) enzyme exerts minor effects on metabolic process (or claims few metabolic requirement). While such explanation for the limited fitness cost by Tet(X6) needs more experimental explorations.

### Inability of tigecycline to treat Tet(X)-producing *E. coli*

Since that Tet(X6) possesses the activity of oxygenating tetracycline (**Fig. S7**) and its derivative tigecycline (**Figs 3-4**), it is reasonable to anticipate it might interfere effectiveness of tigecycline in clinical sector. As very recently Song and coworkers ^45^ established in the case of *mcr-1*, we also adopted the infection model of *Galleria mellonella* (*G. mellonella*) to address this prediction (**Fig. 10A**). Given the constitutive expression of resistance enzymes in the recipient host, both *mcr-1* and *tet*(X6) were fused with the native promoters and then cloned into a low-copy vector of pWSK129 to give pWSK129::P*mcr-1* and pWSK129::P2*tet*(X6), respectively (**Table S1**). Subsequently, these two recombinant plasmids were separately engineered into the well-known virulent strain EDL933 of *E. coli* O157:H7, which generated strain FYJ4039 carrying pWSK129::P*mcr-1* and strain FYJ4040 containing pWSK129::P2*tet*(X6) (**Table S1**). Unlike the negative control group that are consistently killed within 36 hrs after the treatment of PBS alone (**Fig. 10B**), 5 out of 8 larvae survived in the treatment of colistin (7.5mg/kg), 1h post-infection of virulent EDL933 strains (**Fig. 10B**). Notably, all the eight larvae were killed by the MCR-1-producing pathogenic strains of EDL933, regardless of colistin treatment (**Fig. 10B**). This revealed that *mcr-1* renders colistin inefficient in the infection model of *G. mellonella*. In fact, similar scenarios were observed with *mcr-1* in the infection models of *G. mellonella* ^45^ and mouse thighs ^27,45^. As expected, the tigecycline-based therapy (4mg/kg) seemed effective in part (if not all), because that 6 of 8 larvae (75%) are alive within the whole monitoring period of 72hrs post-infection of virulent *E. coli* O157:H7 (**Fig. 10C**). Whereas in the negative control of PBS, none of larvae is exempt from the killing by the virulent strain EDL933 (**Fig. 10C**). Not surprisingly, nearly all the 8 infected *G. mellonella* still were dead, despite that they were treated with tigecycline (**Fig. 10C**). Consistent with that of *tet*(X4) reported by Sun *et al*. ^19^, the observation also enabled us to believe that Tet(X6) abolishes clinical effectiveness of tigecycline. In summary, MCR-1 and Tet(X6) are posing challenges to the renewed interests of colistin and tigecycline, as two last-resort antibiotics used in clinical therapies against severe infections by pathogenic bacteria with multiple resistance.

**Fig. 10.**
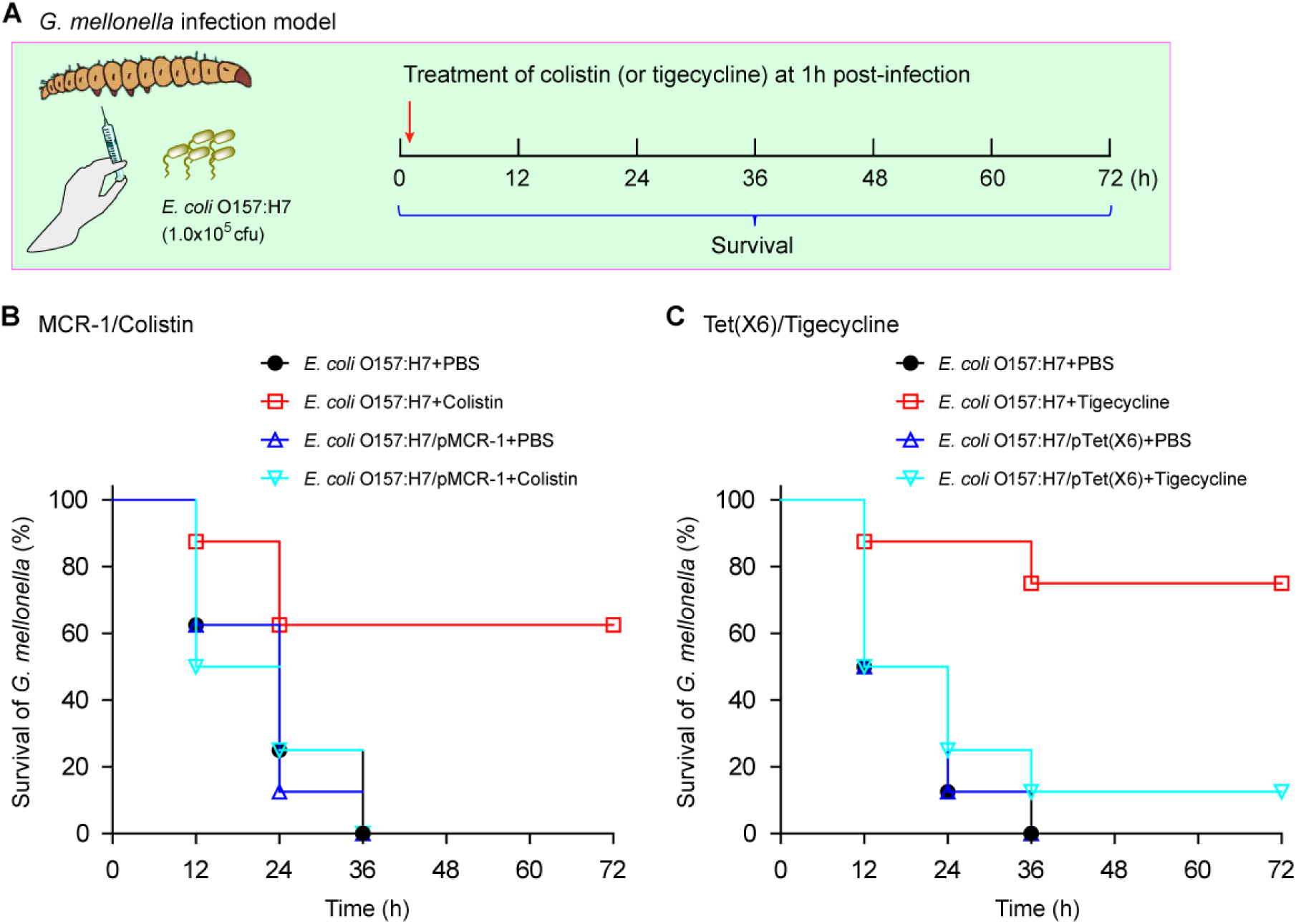
Survival curves of *G. mellonella* suggested that the two treatments of both colistin and tigecycline are ineffective for the *E. coli* infections producing either MCR-1 or Tet(X6) **A.** Schematic representative for *G. mellonella* infection model **B.** The failure in colistin treatment as for *G. mellonella* infected with *mcr-1*-harboring virulent *E. coli* **C.** Tet(X6) renders tigecycline useless in the infection model of *G. mellonella* In addition to the virulent strain EDL933 of EHEC O157:H7, the two derivative strains were included here (**Table S1**). Namely, they referred to FYJ4039 with pWSK129::P*mcr-1*(panel B) and FYJ4040 harboring pWSK129::P2*tet*(X6) (panel C). A representative result is given from three independent assays.

## Conclusions

The pMS8345A, a large IncHI2-type MDR-plasmid is firstly identified by Beatson and coworkers ^24^ to coexist with a big ColV-like virulence plasmid in the ST95 virulent lineage of *E. coli*. This alarms us that the spread of such pathogen might herald an era of post-antibiotic where we stand. The data we report here furthers our understanding tigecycline resistance mechanism of TetX family enzymes (**Fig. 4E**). To the best of knowledge, it is a first report addressing a case of the co-transfer of *tet*(X6) and *mcr-1* by a single plasmid. Since that no known mobile elements are adjacent to *tet*(X6), we hypothesize that the transposon of “IS*Apl1*-*mcr-1-pap2-*IS*Apl1*” mediates the transfer of *mcr-1* into this plasmid. The discovery of Tet(X6), a new member of Tet(X) family, allows us to engineer an array of Tet(X)-expressing bacteria, which paves a way to the development of bioremediation strategy for the environmental tetracycline contamination in the agricultural/industrial productions.

Also, the co-carriage of *tet*(X6) and *mcr-1* on a single IncHI2-type plasmid is far different from the observation by Sun *et al*. ^19^ that the co-existence of *tet*(X4) and *mcr-1* is mediated by two distinct plasmids in an *E. coli* clone. Unlike that the mobility of *tet*(X4) relies on IS*CR2*-mediated transposition ^18^, the gain of *tet*(X6) transferability is not clear (**Fig. 1B**). Not surprisingly, the IncHI2-type plasmid carries *mcr-1* along with *tet*(X6) here, because that it has ever been found to act as a vehicle of global *mcr-1* dissemination ^8,46^. As expected, two types of different antibiotics (colistin and tigecycline) consistently stimulate the formation of hydroxyl radicals in *E. coli* (esp. ROS, in **Fig. 6**), which might constitute an additional example and/or evidence for an improved postuate of “efficient antibiotic killing associated with bacterial metabolic state [ATP ^47-49^ and ROS ^50-53^]. As for a given version of MCR resistance determinants ^39,41,42^, the recipient bacterial host has been demonstrated to give fitness cost exemplified with the delayed growth prior to the entry into log-phase. It is reasonable that the presence of either *mcr-1* or *tet*(X6) also cause metabolic fitness to some extent. Consistent with scenarios with MCR-like members by Yang *et al*. ^38^ and Zhang *et al*. ^39,40,42^, we verified the fitness cost caused by MCR-1 (**Figs 7-8**). This is in part (if not all) explained by the fact that bacterial membrane integrity is altered by MCR-1-mediated lipid A remodeling ^9^. In contrast, the expression of *tet*(X6) does not lead to metabolic burden detected (**Figs 7-8**), which is probably because that Tet(X6) destructs the antibiotic of glycyl-cycline tigecycline, rather than the ribosome target ^10^. It is unusual, but not without any precedent. A similar scenario was seen with the other resistance enzyme β-lactamase-encoding gene *bla*_TEM1b_ (i.e., no fitness cost is correlated with it) ^38^. Therefore, we are not surprised with that no synergistic fitness arises from the co-carriage of *tet*(X6) and *mcr-1* in *E. coli*. Although that a growing body of new *tet*(X) variants [*tet*(X6) to *tet*(X17)] have been proposed in this study (**Fig. 2**), most members of this family, apart from *tet*(X6), await experimental demonstration in the near future. This is because that rare case of cryptic version might occur naturally, such as the prototypical *tet*(X) [we called *tet*(X0)] preexisting in an obligate anaerobe *Bacteroides fragilis* ^21^. Given that i) tigecycline and colistin both are one of few alternative options to combat against carbapenem-resistant Enterobacteriaceae and Acinetobacter species, ii) both MCR-1 ^6^ and Tet(X4) ^19^ have accordingly rendered colistin and tigecycline ineffective in the therapy of mice with MDR infection, the co-occurrence and co-transfer of *tet*(X6) and *mcr-1* by a single plasmid amongst epidemic pathogens is a risky challenge to public health and clinical therapies.

Taken together, it is plausible and urgent to introduce *mcr* variants along with, but not only limited to, *tet*(X) variants in the routine national (and/or international) investigation in the context of “one health” (environmental/animal/human sectors). Along with major findings of other research group ^12^, functional definition of Tet(X3) and its homologue Tet(X6) here extends mechanistic insights into Tet(X) tigecycline resistance, and even benefit the development of anti-Tet(X) resistance enzyme inhibitors.

## Materials and Methods

### Sequencing, assembly and annotation of plasmids

The plasmid pDJB-3 was isolated from the colistin-resistant *E. coli* DJB-3 of swine origin, verified with *mcr-1*-specific PCR, and then subjected to genome sequencing with the Hiseq X ten PE150 sequencer platform (Illumina, USA). As a result, the DNA library of pDJB-3 plasmid prepared by KAPA Hyper Prep Kit (Roche, Basel, Switzerland) gave a pool of 150 bp paired-end reads that are destined to be assembled into a contig by the SPAdes Genome Assembler (version 3.11.0). A BLASTN search was conducted to probe whether or not the resultant *mcr-1*-containing contig has a best-hit plasmid candidate. Together with Sanger sequencing, PCR was applied to close all the suspected gaps.

The resultant plasmid genome was annotated through the prediction of open reading frames (ORFs) with RAST (rapid annotation using subsystem technology, http://rast.nmpdr.org). PlasmidFinder 1.3 (https://cge.cbs.dtu.dk/services/PlasmidFinder-1.3/) was used to type the plasmid incompatibility, and ResFinder 3.1 (https://cge.cbs.dtu.dk/services/ResFinder/) was applied to screen possible antimicrobial resistance genes. The plasmid map was given with GenomeVx (http://wolfe.ucd.ie/GenomeVx/), and its linear alignment was proceeded with Easyfig ^54^.

### Plasmid conjugation experiments

As recently described by Sun *et al*. ^27^, the experiments of plasmid conjugation were performed, in which the rifampin-resistant *E. coli* recipient strain EC600 (and/or strain DJB-3) acted as a donor. In brief, overnight cultures were re-grew in LB broth, donor and recipient strains were mixed at the logarithmic phase and spotted on a filter membrane, and then incubated at 37°C overnight. Subsequently, bacteria were washed from filter membrane and spotted on LB agar plate containing 400µg/ml rifampin and 4 µg/ml colistin for selection of transconjugants. The suspected transformants were validated with PCR assays.

### Molecular and microbial manipulations

With all the known *tet*(X) variants [*tet*(X0)-*tet*(X5)] as queries, BLASTN was carried out. In particular, a *tet*(X3)-based search returned a plasmid pMS8345A with significant hit, leading to the discovery of new variant of *tet*(X6). Then, *tet*(X6) was synthesized *in vitro*, and cloned into pABD24, giving pABD24::*tet*(X6) (**Table S1**). Following the verification of its identity with direct DNA sequencing, this recombinant plasmid was introduced into the MG1655 strain of *E. coli* to assess its role *in vivo*. The generation of all the point-mutants of *tet*(X6) were based on pBAD24::*tet*(X6) (**Table S1**) using site-directed mutagenesis kit (Vazyme Biotech), along with an array of specific primers (**Table S2**). To test relationship of MCR-1 with Tet(X6), *mcr-1* was cloned into pWSK129, giving pWSK::*mcr-1*, compatible with pBAD::*tet*(X6) within a single *E. coli* colony (**Table S1**). After experimental validations of phenotypic clolistin resistance (and/or tigecycline resistance), all the bacterial were subjected to routine isolation of crude lipo-polysaccharides-lipid A as earlier recommended by Caroff *et al*. ^55^. The identity of purified lipid A species were verified with MALDI-TOF/TOF mass spectrometry (Bruker UltrafleXtreme, Germany) ^25^.

As recently described with *tet*(X4) ^20^, the ability of *tet*(X6) and *tet*(X3) in phenotypic tigecycline resistance was evaluated with LB agar plates containing tigecycline in a series of dilution. The strain expressing *tet*(X4) is used as a positive control. In addition, the MCR-1 colistin resistance was also judged as we earlier conducted with *mcr-1* ^56^ with little change. All the examined *E. coli* strains were cultivated at 37°C overnight. Overnight cultures were standardized to OD_600_ 0.05, inoculated (1:10; v/v) into 96-well glass-bottomed plates in fresh LB broth ± arabinose or lactose (0, 0.02%, and 0.2%, w/v), and shaken (180 r.p.m) at 37°C. Of note, arabinose acted as an inducer of pBAD24, and lactose was used to trigger expression of pWSK129-based MCR-1. As the establishment with NMCR-1 ^39^ and MCR-3/5 ^41,42^, bacterial growth curves were automatically plotted with spectrophotometer (Spectrum lab S32A) to evaluate the fitness cost caused by *tet*(X6) and *mcr-1*. During the total period of 20 hours, the value of optical absorbance (i.e., OD600) was consistently recorded at an interval of 1 hour.

### Bioassays for tigecycline destruction

The hydrolytic activity of Tet(X6) [and/or Tet(X3)] enzyme was determined as Balouiri *et al*. ^57^ described with little change. In brief, the strain of MG1655 harboring pBAD24::*tet*(X6) [or pBAD24::*tet*(X3)] was cultivated overnight on LB agar plates supplemented with 0.1% arabinose. As a result, bacterial colonies stripped, were suspended with 0.5ml of LB broth containing 0.1% arabinose and 2.5mg/ml tigecycline, whose optical density at 600nm (OD600) was adjusted to about 2.0. Then, the suspension cultures were proceeded to 8h stationary growth at 37°C. Following centrifugation (13,600rpm, 20min) and filtration (at 0.22μm cut-off), bacterial supernatants were prepared. The *E. coli* DH5α here referred to an indicator strain of tigecycline susceptibility. Of note, the overnight culture of *E. coli* DH5α (∼100μl) was spread on a LB agar plate, which is centered with a paper disk of 6mm diameter. To visualize the inhibition zones, the supernatant of interest (∼20µl) was spotted the paper disk on the aforementioned bioassay plates, and incubated at 37°C for 16h. The negative-control denotes the supernatant from *E. coli* MG1655 bearing the empty vector pBAD24 (**Table S1**), and the blank control referred to the LB broth containing 2.5mg/ml tigecycline.

### Expression, purification and identification of Tet(X) enzymes

To produce the Tet(X6) protein and its homologue Tet(X3), the strains of *E. coli* BL21 carrying pET21::*tet*(X6) [and pET21::*tet*(X3)] were engineered (**Table S1**) for the inducible expression via the addition of 0.5 mM isopropyl β-D-1-thiogalactopyranoside (IPTG). Bacterial lysates obtained by a French Press (JN-Mini, China), were subjected to 1h of centrifugation at 16,800 rpm at 4°C, and the resultant supernatants were incubated with pre-equilibrated Ni-NTA agarose beads on ice for 3 hours. Following the removal of protein contaminants, the Tet(X6) [and/or Tet(X3)] protein was eluted from the Ni-NTA agarose beads using the elution buffer [20mM Tris-HCl (pH 8.0), 150mM NaCl, 20mM imidazole, and 5%glycerol], and concentrated with a 30kDa cut-off ultra-filter (Millipore, USA). Subsequently, gel filtration was performed to probe solution structure of Tet(X6) [Tet(X3)], using a Superdex 200/300GL size exclusion column (GE Healthcare). The purity of protein pooled from the target peak was judged with SDS-PAGE (15%), and its identity was validated with MALDI-TOF/TOF mass spectrometry (LTQ orbitrap Elite, Thermo Fisher).

### Enzymatic activity for Tet(X6) *in vitro*

To confirm the enzymatic activity of Tet(X6) enzyme, the *in vitro* reaction system was established as recently described by Sun *et al*. ^19^ with little change. The components of this assay (50μl in total) consisted of 20mM Tris (pH7.5), 150mM NaCl, 1mM NADPH, 4mg/ml tigecycline, and the purified enzyme [2mg/ml for either Tet(X6) or Tet(X3)]. Following the maintenance (∼12h) of enzymatic reaction at room temperature, the resultant reaction mixture was subjected to further analysis of liquid chromatography mass spectrometry (LC/MS) using an Agilent 6460 triple quadrupole mass spectrometer (Agilent Technologies, USA) ^18^. As for LC/MS here, it was carried out as follows: i) Nitrogen acted as the sheath gas and drying gas, the nebulizer pressure was set to 45 psi, and the flow rate of drying gas was 5 liter/min. The flow rate and temperature of the sheath gas were 11 liter/min and 350°C, respectively; ii) Chromatographic separation proceeded on a Zorbax SB C8 column (150 × 2.1mm, 3.5µm); iii) Mass spectrometric detection was completed using an electrospray ionization (ESI) source in positive mode. Scan range was 100∼1000amu; and iv) The resultant data was processed with Agilent Mass Hunter Workstation.

### The steady state kinetic assay of Tet(X6)

The decrease in absorbance corresponding to tigecycline hydroxylation by Tet(X6) were monitored at 400nm (ε_400_ =4300 M^−1^cm^−1^) over 6 min. To determine the steady-state kinetics parameters for Tet(X6), we measured initial velocities (*V*_0_) of tigecycline inactivation at varied concentration of tigecycline (60μM, 80μM, 100μM, and 120μM), 1mM NADPH, 5mM MgCl_2_, 0.5μM Tet(X6) protein, 20mM Tris-HCl (pH8.5) concentrations at 37°C ^15^. Each 200μl reaction in 96-well micro-titre was monitored using SPECTROstar ^Nano^. All assays were performed in triplicate. Steady-state kinetic parameters were determined by fitting initial reaction rates (*V*_0_). The data was analyzed according to the standard Michaelis-Menten equation. The double-reciprocal plot featuring with the formula “1/V_0_=(Km/V_max_)/[S]+1/V_max_” was used to calculate the *K*_m_. Accordingly, *V*_0_=*V*_max_ [S]/(*K*_m_+[S]). The catalytic constant *k*_cat_ was determined according to the *V*_max_ = *k*_cat_ [E_0_], and E_0_ is total enzyme concentration ^58^.

### Flow cytometry

Mid-log phase cultures (OD600, ∼1.0) were prepared for the detection of intra-cellular reactive oxygen species (ROS). The oxidant sensor dye, DCFH2-DA (sigma) was mixed with bacterial strains and kept for 0.5h. Accordingly, the 2.0mg/ml of antibiotics (colistin and/or tigecycline) were supplemented. Then, bacterial samples (10^5^∼10^6^) diluted with 0.85% saline were subjected to the analysis of flow cytometry ^40,42^. The resultant FACS data was recorded with a BD FACSVerse flow cytometer through counting 10,000 cells at a flow rate of 35ml/min (and/or 14ml/min). In particular, DCFH florescence was excited with a 488nm argon laser and emission was detected with the FL1 emission filter at 525nm using FL1 photomultiplier tub.

### Confocal microscopy

As recently described ^38^, confocal microscopy was conducted to examine the potential effects on bacterial viability exerted by resistance enzymes [MCR-1 and/or Tet(X6)]. Prior to assays of confocal microscopy, mid-log phase cultures were processed with the LIVE/DEAD BacLight™ Bacterial Viability Kit (Cat. No. L7012) ^38^. Namely, the three strains tested here included i) *E. coli* MG1655 (*mcr-1*/pWSK), ii) MG1655 [*tet*(X6)/pBAD], and iii) MG1655 [*tet*(X6)/pBAD and *mcr-1*/pWSK). Of note, 0.2% lactose is an inducer of *mcr-1* expression, and 0.2% arabinose acts as an activator for Tet(X6) enzyme production. After the removal of supernatants, bacterial biofilms were stained with 3% LIVE/DEAD kit solution, and maintained at room temperature in the dark for 15 minutes. Photographs were captured by the confocal laser scanning microscopy (Zeiss LSM 800) with a 63× oil immersion lens and analyzed using COMSTAT image analysis software. The Tukey–Kramer multiple comparison post hoc test was applied to judge the COMSTAT data. Statistical significance was set at p< 0.01 with T-test.

### Infection model of *G. mellonella*

To probe possible interferences of *mcr-1* and/or *tet*(X6) in the anti-bacterial treatment with colistin (and/or tigecycline), the infection model of *Galleria mellonella* (*G. mellonella*) was applied here. Prior to bacterial infections, the larvae of *G. mellonella* (Tianjin Huiyude Biotech Company, Tianjin, China) was assessed as for the weight (0.3-0.4g each) and its active status, and then grouped appropriately (8 per group). The mid-log phase cultures of the virulent *E. coli* (EHEC O157:H7) with or without plasmid-borne *mcr-1* [and/or *tet*(X6)] were prepared (**Table S1**), and then suspended with 1xPBS buffer, in which the final OD600 is 0.1. As recently Song *et al*. performed ^45^ with minor change, each larvae was injected with 10ul of bacterial solution (1.0x 10^5^ cfu) at the left posterior gastropoda. After 1h post-challenge, the infected larvae separately received the different treatments on the right posterior gastropoda ^45^. Namely, they referred to PBS, colistin (7.5mg/kg), and tigecycline (4mg/kg) ^45^. Survival rate of *G. mellonella* was monitored over 72hrs, of which an interval is 12hrs. Three biological replicates were performed.

### Bioinformatics

Multiple sequence alignments of Tet(X) variants at the levels of both amino acids and nucleic acids proceeded with ClustalOmega (https://www.ebi.ac.uk/Tools/msa/clustalo). Consequently, the phylogeny of Tet(X) was generated with TreeView (https://www.treeview.co.uk/). Tet(X6) was structurally modeled using Swiss-Model (https://swissmodel.expasy.org/interactive) ^59^, in which the structural template detected refers to Tet(X2) (PDB: 2Y6Q) ^16,58^. Both GMQE (global model quality estimation) and QMEAN (a global and local absolute quality estimate on the modeled structure) was applied to judge the quality of the modeled structure. Finally, structural presentation and cavity illustration of Tet(X6) was given with PyMol (https://pymol.org/2).

### Accession numbers

Nucleotide sequence data of *tet*(X6) reported here is available in the Third-Party Annotation Section of the DDBJ/ENA/GenBank databases under the accession number TPA: BK011183. The full genome sequence of the *mcr-1*-harboring plasmid pDJB-3 of swine origin is accessed under the accession number: MK574666.

## Acknowledgements

We would like to thank Dr. Yuanyuan Zhang for the technical assistance in flow cytometer assays with BD FACSVerse (Shared Management Platform for Large Instrument, College of Animal Sciences, Zhejiang University). We are grateful to Prof. Jian-hua Liu (South China Agricultural University, Guangzhou, China) for providing us the *mcr-1*-harboring plasmid, pHNSHP45-2. This work was supported by National Key R&D Program of China (2017YFD0500202, YF) and National Natural Science Foundation of China (31830001, 31570027 & 81772142, YF). Dr. Feng is a recipient of the national “Young 1000 Talents” Award of China.

## Author contributions

YF designed and supervised this study; YF, YX, LL and HZ conducted experiments and analyzed data; YF and HZ contributed regents and interpreted data; YF and HZ drafted and reviewed this manuscript.

## Competing Interests

We declare that no conflict of interest is present.

## Supporting information

### Supporting tables

**Table S1.** Bacterial strains and plasmids used in this study

| Strains/plasmids | Description | Origins |
| --- | --- | --- |
| **Strains** |  |  |
| DH5α | A cloning host of *E. coli* | Lab stock |
| MG1655 | A wild-type strain of *E. coli* K-12 | Lab stock |
| EDL933 | A virulent strain of *E. coli* O157:H7 | Lab stock |
| FYJ796 | MG1655 carrying pBAD24 | Lab stock |
| FYJ4000 | MG1655 carrying pBAD24::tet(X6) | Lab stock |
| FYJ4019 | MG1655 carrying pBAD24::tet(X6) | This work |
| FYJ4020 | MG1655 carrying pBAD24::tet(X3) | This work |
| FYJ4021 | MG1655 carrying pWSK129::mcr-1 | This work |
| FYJ4022 | MG1655 carrying pWSK129::mcr-1 and pBAD24::tet(X6) | This work |
| FYJ4023 | BL21 carrying pET21a::tet(X6) | This work |
| FYJ4024 | MG1655 carrying pBAD24::tet(X6) (E36A) | This work |
| FYJ4025 | MG1655 carrying pBAD24::tet(X6) (R37A) | This work |
| FYJ4026 | MG1655 carrying pBAD24::tet(X6) (R107A) | This work |
| FYJ4027 | MG1655 carrying pBAD24::tet(X6) (Q182A) | This work |
| FYJ4028 | MG1655 carrying pBAD24::tet(X6) (R203A) | This work |
| FYJ4029 | MG1655 carrying pBAD24::tet(X6) (H224A) | This work |
| FYJ4030 | MG1655 carrying pBAD24::tet(X6) (G226A) | This work |
| FYJ4031 | MG1655 carrying pBAD24::tet(X6) (D301A) | This work |
| FYJ4032 | MG1655 carrying pBAD24::tet(X6) (P308A) | This work |
| FYJ4033 | MG1655 carrying pBAD24::tet(X6) (Q312A) | This work |
| FYJ4034 | MG1655 carrying pBAD24::tet(X6) (M365A) | This work |
| FYJ4035 | BL21 carrying pET21a::tet(X3) | This work |
| FYJ4036 | MG1655 carrying pWSK129::P1 tet(X6) | This work |
| FYJ4037 | MG1655 carrying pWSK129::P2 tet(X6) | This work |
| FYJ4038 | EDL933 carrying pWSK129 | This work |
| FYJ4039 | EDL933 carrying pWSK129::Pmcr-1 | This work |
| FYJ4040 | EDL933 carrying pWSK129::P2 tet(X6) | This work |
| **Plasmids** |  |  |
| pBAD24 | The arabinose-inducible expression vector, Amp<sup>R</sup> | Lab stock |
| pWSK129 | The lactose-activated expression vector, Km<sup>R</sup> | Lab stock |
| pHNSHP45-2 | A big *mcr-1*-bearing plasmid (251,493bp) from the *E. coli* SH45 strain (acc. no.: KU341381) | 29 |
| pBAD24::tet(X4) | A pBAD24 carrying tet(X4) at the two cuts of EcoRI and Sall, Amp<sup>R</sup> | 20 |
| pBAD24::*tet*(X6) | A pBAD24 carrying *tet*(X6) at the two cuts of EcoRI and Sall, Amp<sup>R</sup> | This work |
| pBAD24::*tet*(X3) | A pBAD24 carrying *tet*(X3) at the two cuts of EcoRI and Sall, Amp<sup>R</sup> | This work |
| pWSK129::*mcr-1* | A pWSK129 carrying *mcr-1* at the two cuts of Sall and EcoRI, Km<sup>R</sup> | This work |
| pET21a::*tet*(X6) | A pET21a vector carrying *tet*(X6) at the two cuts of NdeI and XhoI, Amp<sup>R</sup> | This work |
| pET21a::*tet*(X3) | A pET21a vector carrying *tet*(X3) at the two cuts of NdeI and XhoI, Amp<sup>R</sup> | This work |
| pBAD24::*tet*(X6) (E36A) | A derivative of pBAD24 encoding the point-mutant E36A of *tet*(X6), Amp<sup>R</sup> | This work |
| pBAD24::*tet*(X6) (R37A) | A derivative of pBAD24 encoding the R37A substitution version of *tet*(X6), Amp<sup>R</sup> | This work |
| pBAD24::*tet*(X6) (R107A) | A derivative of pBAD24 encoding the point-mutant (R107A) of *tet*(X6), Amp<sup>R</sup> | This work |
| pBAD24::*tet*(X6) (Q182A) | pBAD24 encoding the Q182A mutant of *tet*(X6), Amp<sup>R</sup> | This work |
| pBAD24::*tet*(X6) (R203A) | pBAD24 encoding the R203A mutant of *tet*(X6), Amp<sup>R</sup> | This work |
| pBAD24::*tet*(X6) (H224A) | pBAD24 encoding the H224A mutant of *tet*(X6), Amp<sup>R</sup> | This work |
| pBAD24::*tet*(X6) (G226A) | pBAD24 encoding the G226A mutant of *tet*(X6), Amp<sup>R</sup> | This work |
| pBAD24::*tet*(X6) (D301A) | pBAD24 encoding the D301A mutant of *tet*(X6), Amp<sup>R</sup> | This work |
| pBAD24::*tet*(X6) (P308A) | pBAD24 encoding the P308A mutant of *tet*(X6), Amp<sup>R</sup> | This work |
| pBAD24::*tet*(X6) (Q312A) | pBAD24 encoding the Q312A mutant of *tet*(X6), Amp<sup>R</sup> | This work |
| pBAD24::*tet*(X6) (M365A) | pBAD24 encoding the M365A mutant of *tet*(X6), Amp<sup>R</sup> | This work |
| pWSK129::*P1 tet*(X6) | pWSK129 carrying the promoter1-linked *tet*(X6) at the two cuts of Sall and EcoRI, Km<sup>R</sup> | This work |
| pWSK129::*P1 tet*(X6) | A derivative of pWSK129 carrying the promoter2-fused *tet*(X6) at the two cuts of Sall and EcoRI, Km<sup>R</sup> | This work |
| pWSK129::*Pmcr-1* | A derivative of pWSK129, of which the two cuts of Sall and EcoRI, is inserted by *mcr-1* along with its native promoter, Km<sup>R</sup> | This work |

**Table S2.**
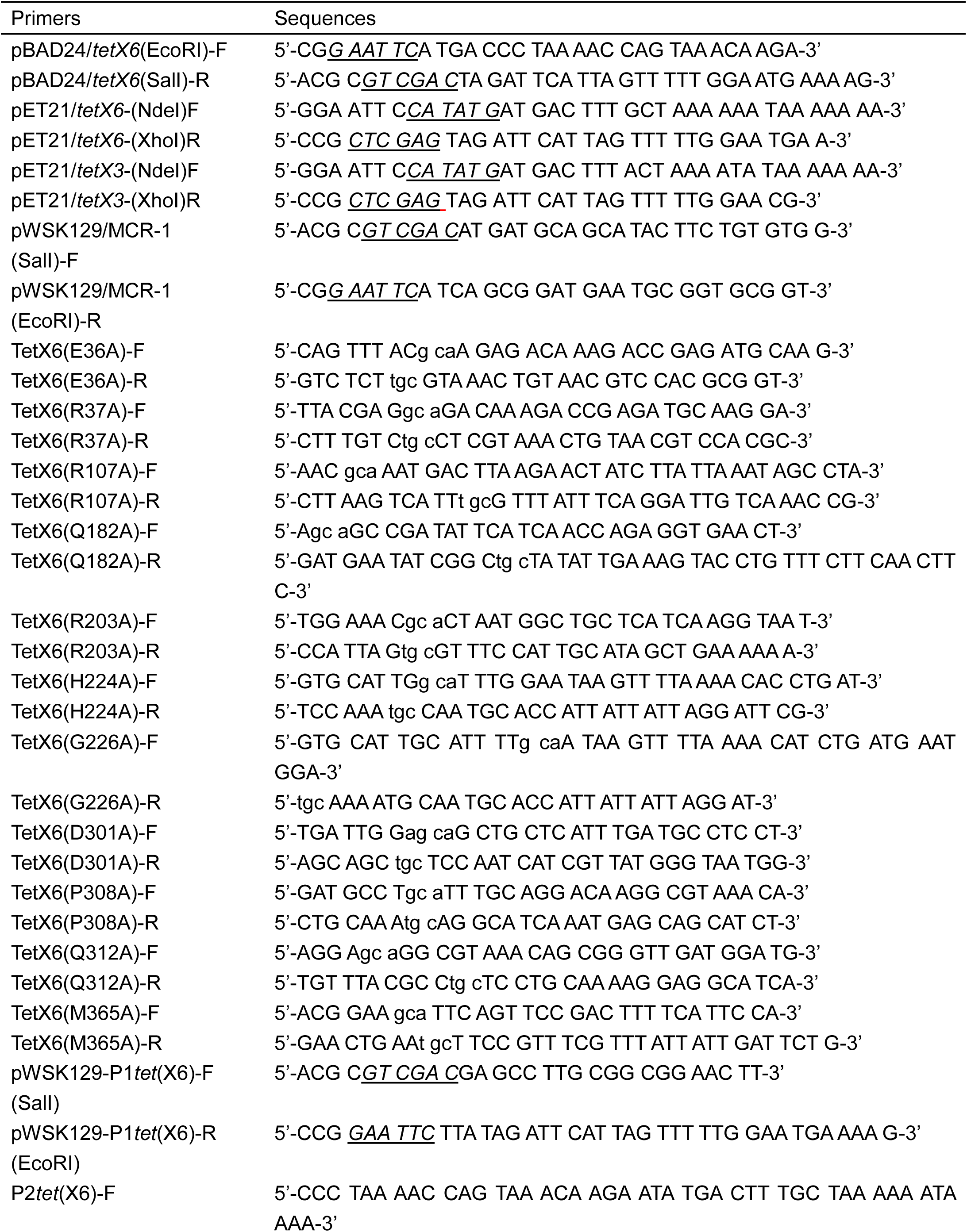
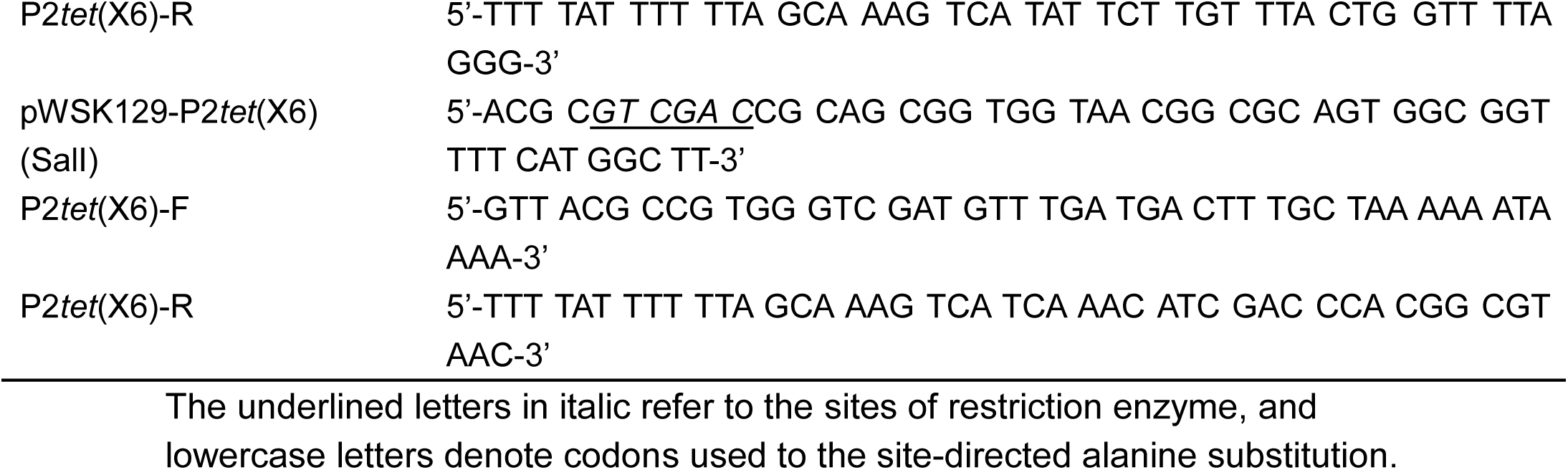
Primers used in this study

### Supporting figures

**Fig. S1.**
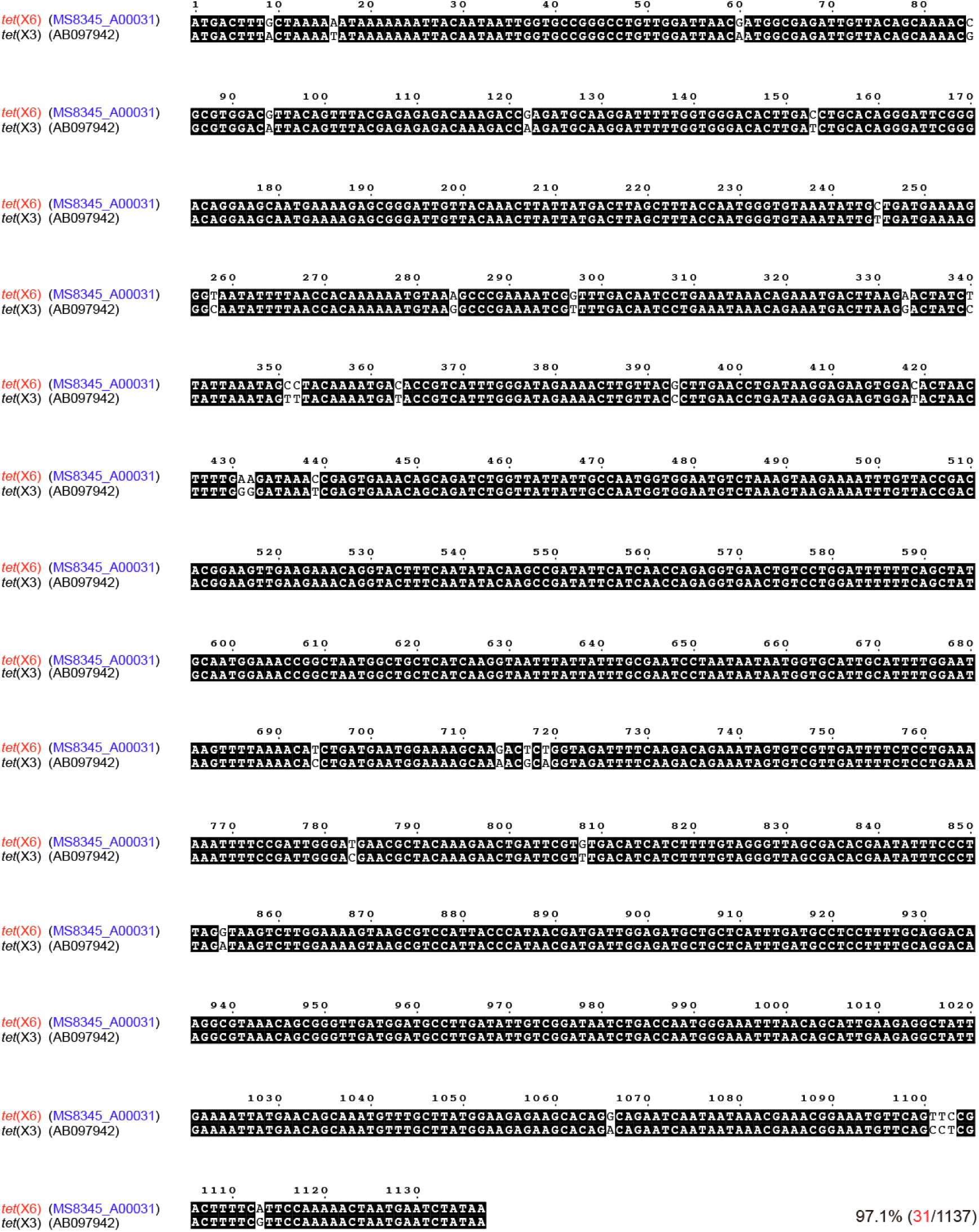
Nucleotide acid sequence analysis of *tet*(X3) and *tet*(X6) To determine the similarity between *tet*(X3) and *tet*(X6), their nucleotide acid sequences were subjected to Clustal Omega (https://www.ebi.ac.uk/Tools/msa/clustalo/), and resultant form of sequence is given with ESPript 3.0 (http://espript.ibcp.fr/ESPript/cgi-bin/ESPript.cgi). Identical residues are in white letters with black background, and different residues are black letters with white background. The identity between *tet*(X3) and *tet*(X6) is 97.1%, with 31 substitution of 1137 residues in total.

**Fig. S2.**
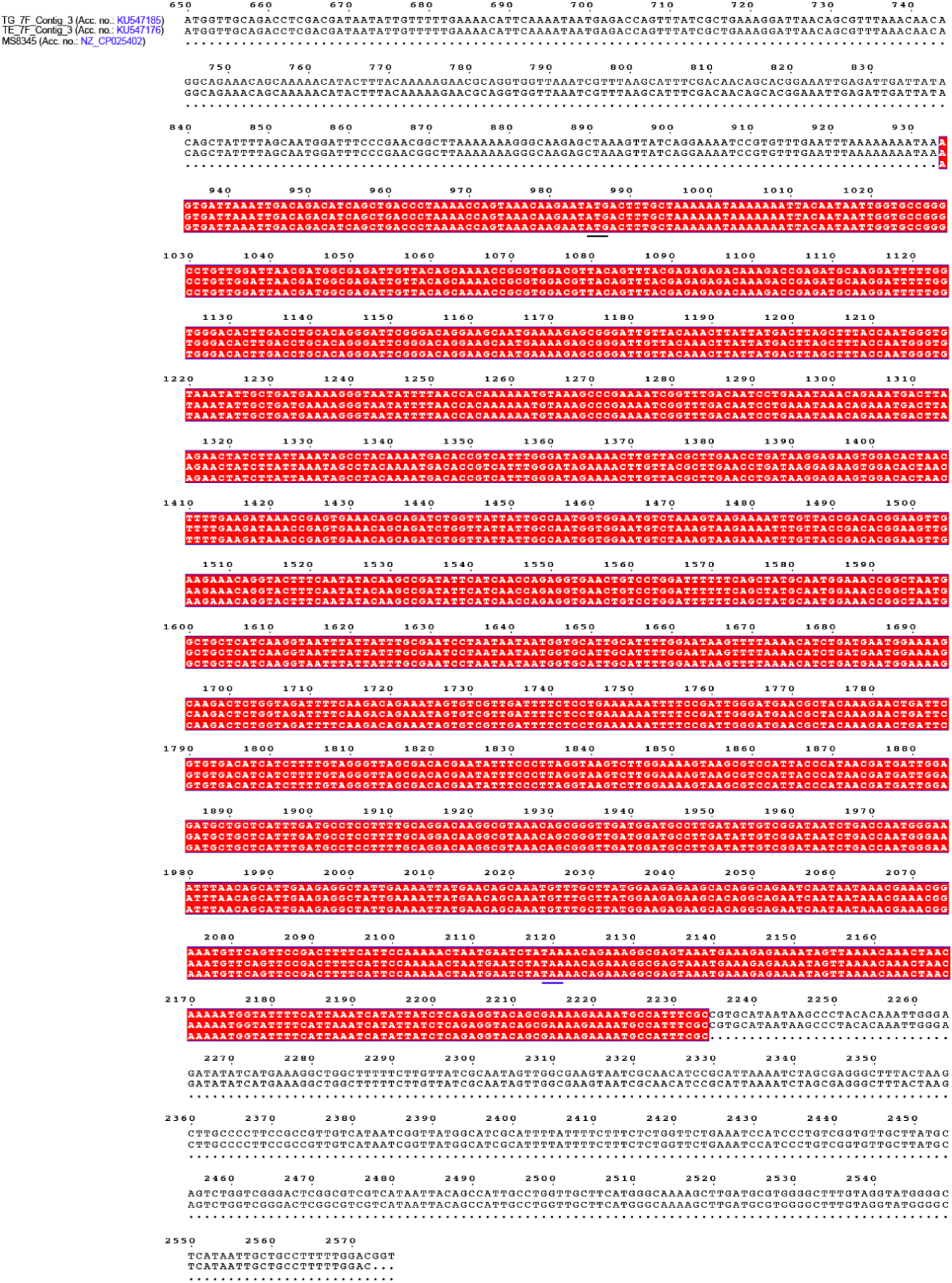
Evidence that an intact *tet*(X6) is only harbored on one plasmid and two contigs. Sequence alignment was conducted as described in **Fig. S1**. An initial codon “ATG” and the stop codon “TAA” are underlined.

**Fig. S3.**
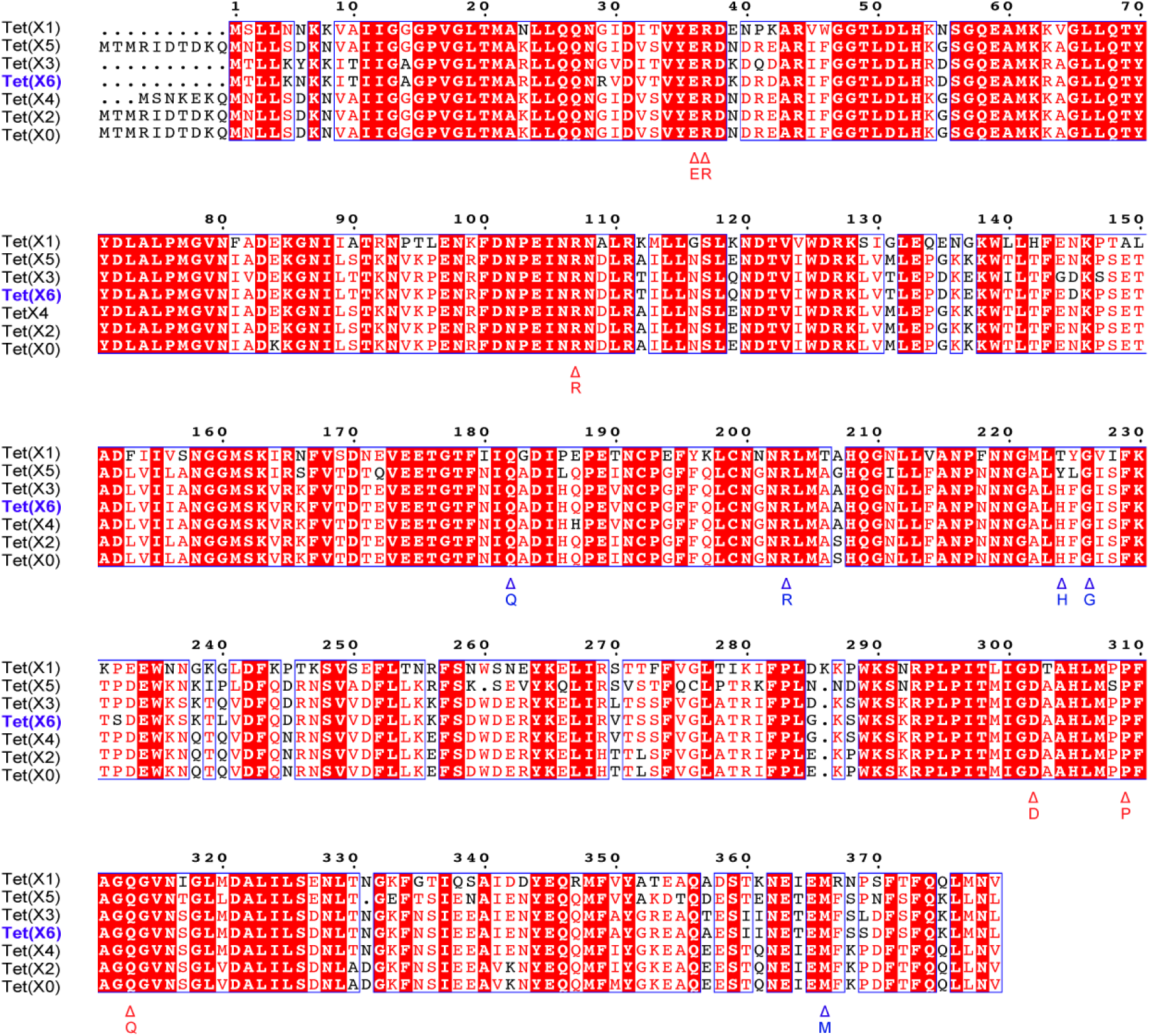
Sequence alignment of the Tet(X6) enzyme with other five homologs (X0 to X5) Multiple sequence alignment was conducted with Clustal Omega (https://www.ebi.ac.uk/Tools/msa/clustalo/), generating its output with ESPript 3.0 (http://espript.ibcp.fr/ESPript/cgi-bin/ESPript.cgi). The putative substrate-loading cavity is composed of FAD-interactive residues (with red triangles) and Tigecycline-binding residues (with blue triangles).

**Fig. S4.**
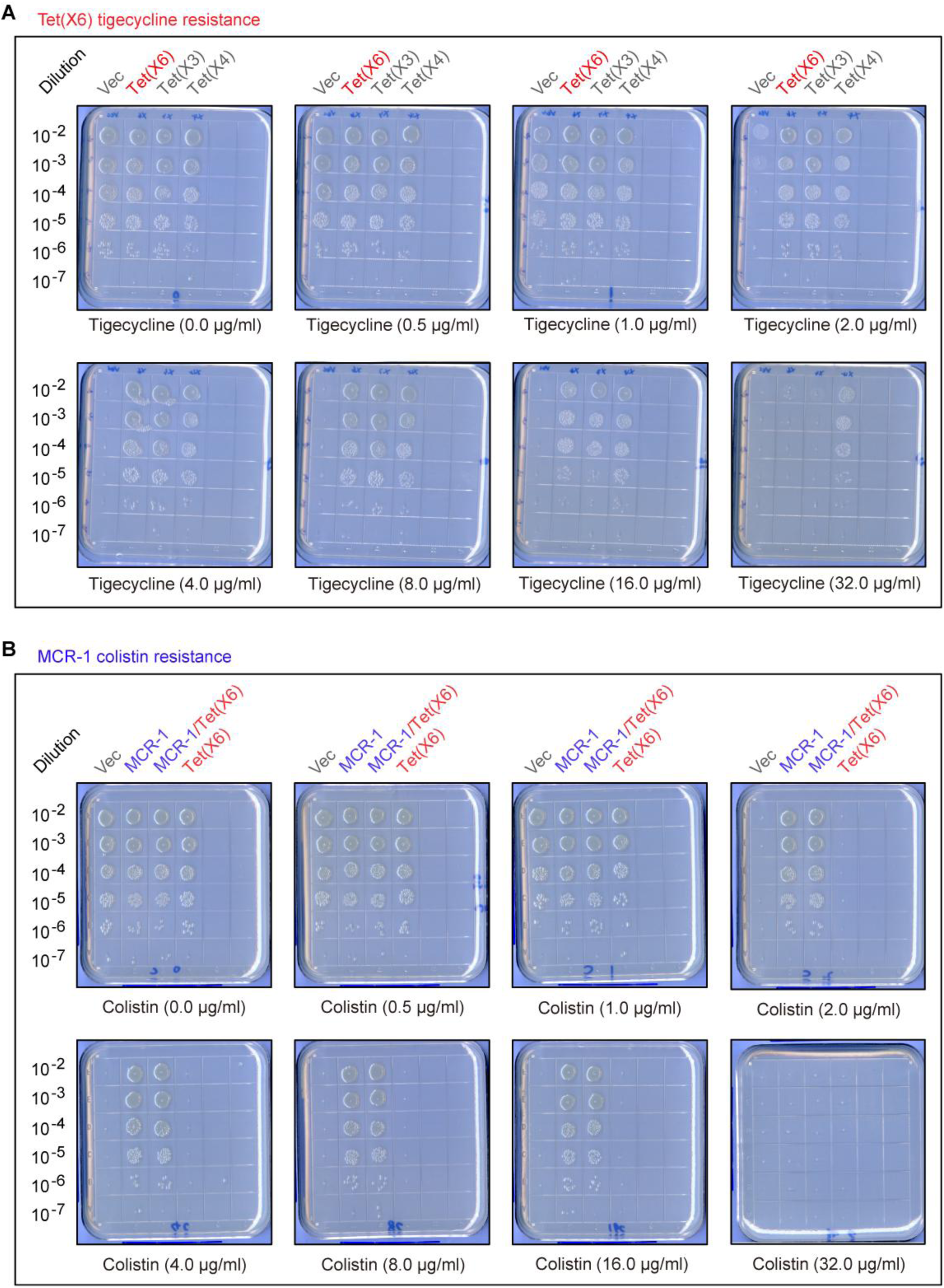
Co-occurrence of *tet*(X6) and *mcr-1* gives co-resistance to colistin and tigecycline. **A.** Contrasting the level of tigecycline resistance by Tet(X) variants The *E. coli* MG1655 carrying different variants of *tet*(X) [X3, X4, and X6] were maintained at 37°C on LB agar plates containing tigecycline in series of dilution. **B.** The MCR-1 confers phenotypic colistin resistance in the *E. coli* MG1655 The derivatives of *E. coli* MG1655 bear *mcr-1* or *tet*(X6) alone (or both) were spotted on LB agar plates containing colistin in series of dilution, and maintained at 37°C for overnight.

**Fig. S5.**
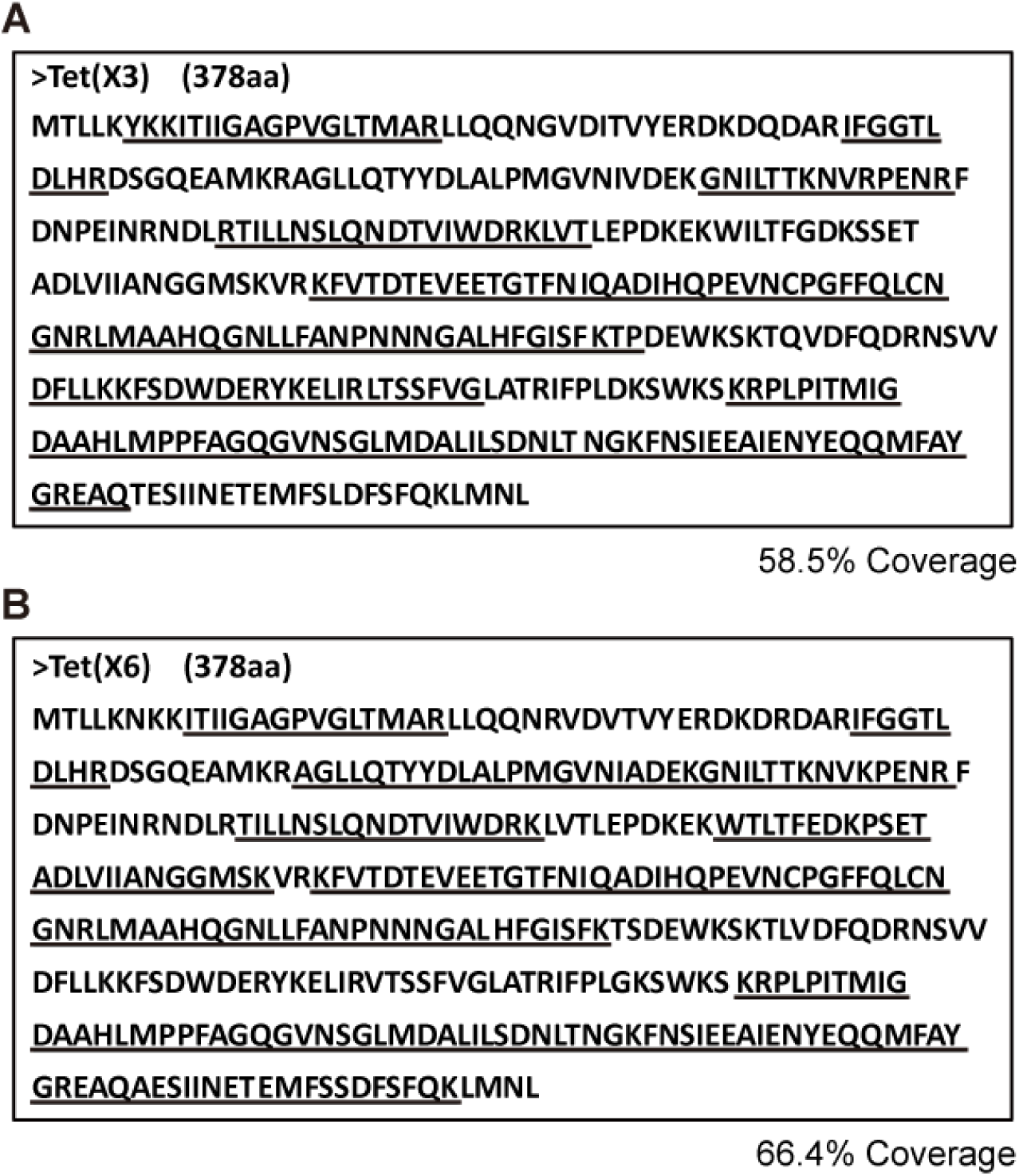
MS identity of the purified two proteins Tet(X3) and Tet(X6) **A.** MS-based identification of polypeptide fragments from the purified Tet(X3) protein **B.** MS-based determination of the purified Tet(X6) protein The underlined letters denote the polypeptides that match Tet(X3) [and/or Tet(X6)] protein.

**Fig. S6.**
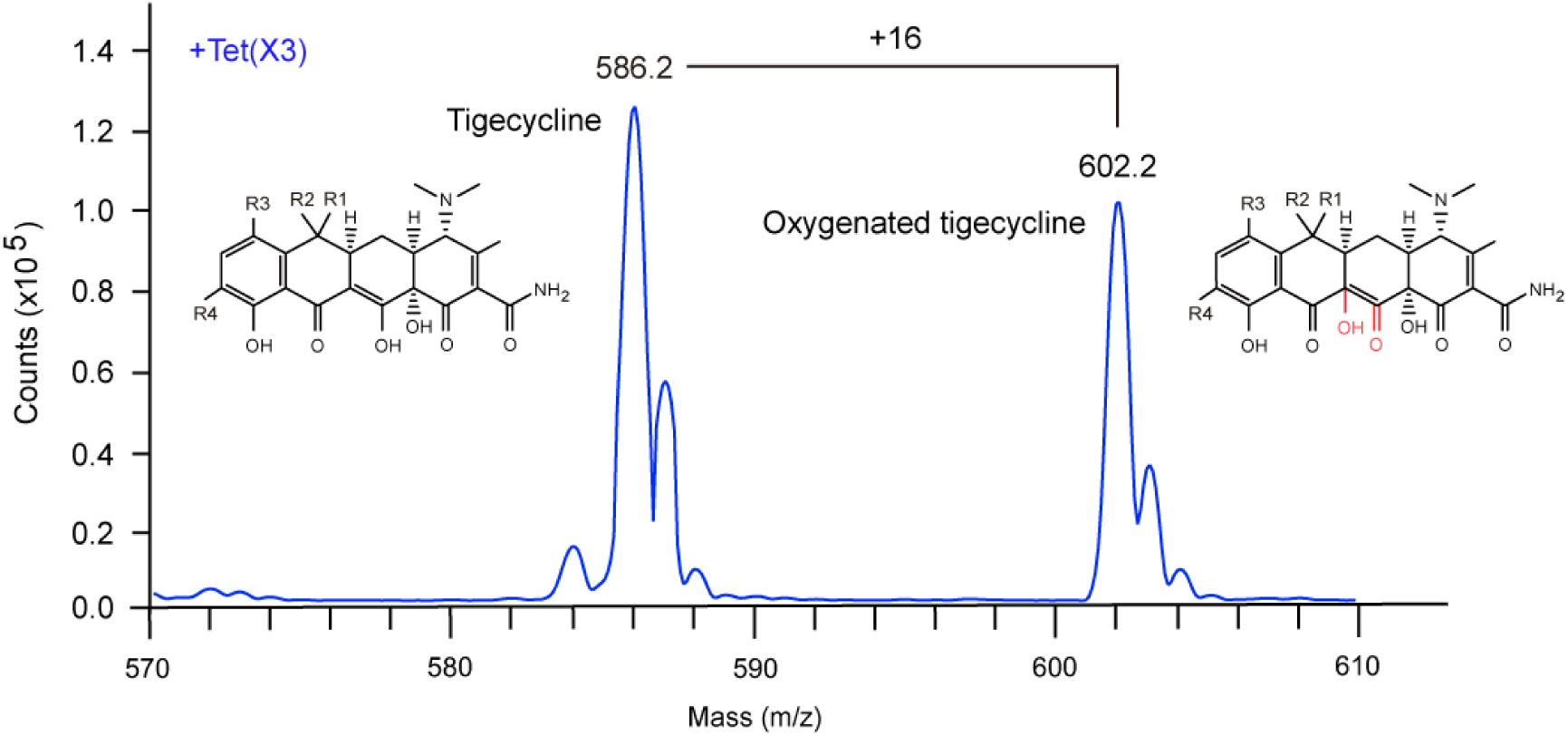
Use of LC/MS to identify the oxygenated product of tigecycline by Tet(X3) enzyme. In the spectrum, the peak of 586.2 (m/z) refers to tigecycline, whereas the other peak of 602.2 (m/z) denotes an oxygenated product of tigecycline. Chemical structures were given with ChemDraw.

**Fig. S7.**
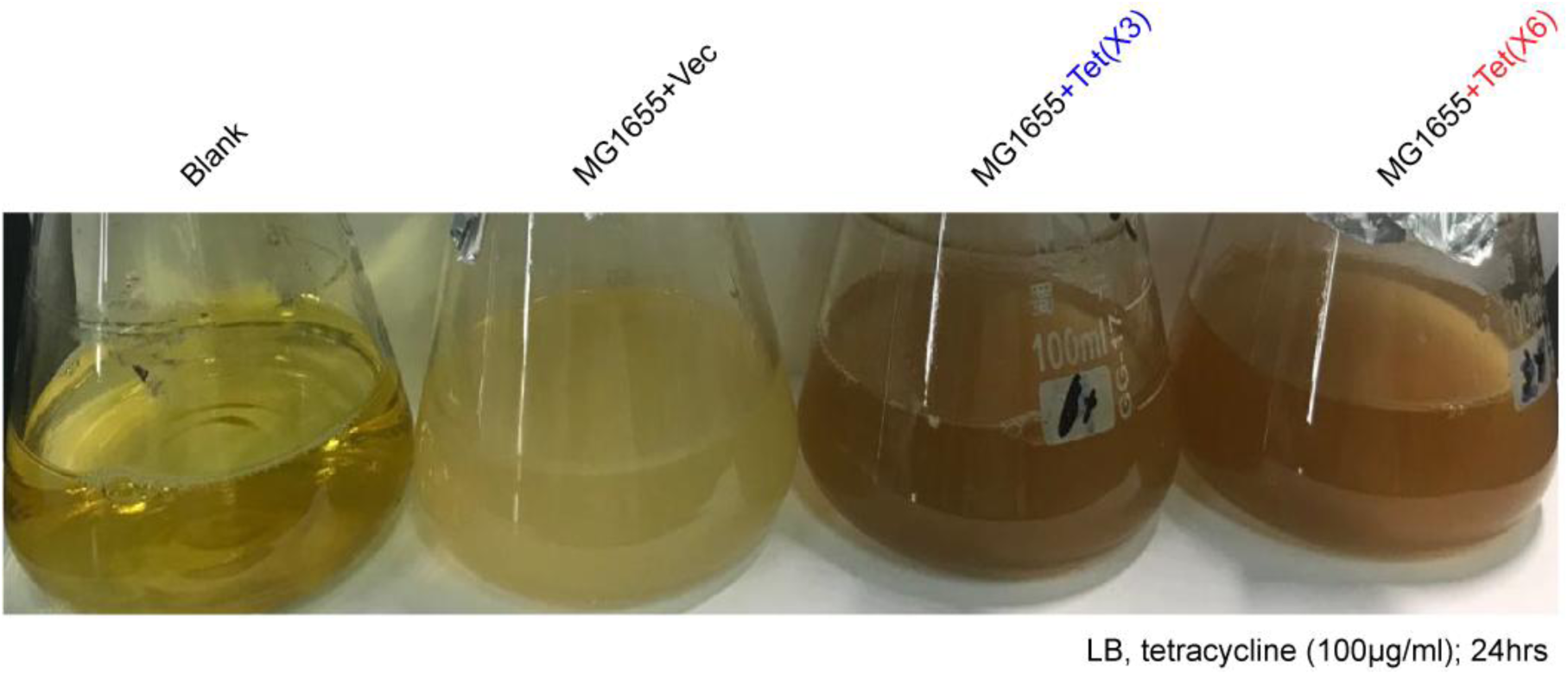
Visualization for destruction of tetracycline by Tet(X3) [and Tet(X6)] enzyme. Unlike the fact that the blank and the negative control (liquid culture of *E. coli* with empty vector alone) display yellow, bacterial culture of Tet(X3) [and/or Tet(X6)]-expressing *E. coli* gives dark. This indicates that the expression of Tet(X3) [and Tet(X6)] leads to the oxygenation of tetracycline, which is fully consistent with the observation of other tetracycline-inactivating enzymes by Forsberg and coworkers ^17^. 0.1% arabinose is used to induce the expression of pBAD24-borne *tet*(X3) [and *tet*(X6)]. Designation: Blank, the LB liquid medium containing tetracycline; Vec, pBAD24

**Fig. S8.**
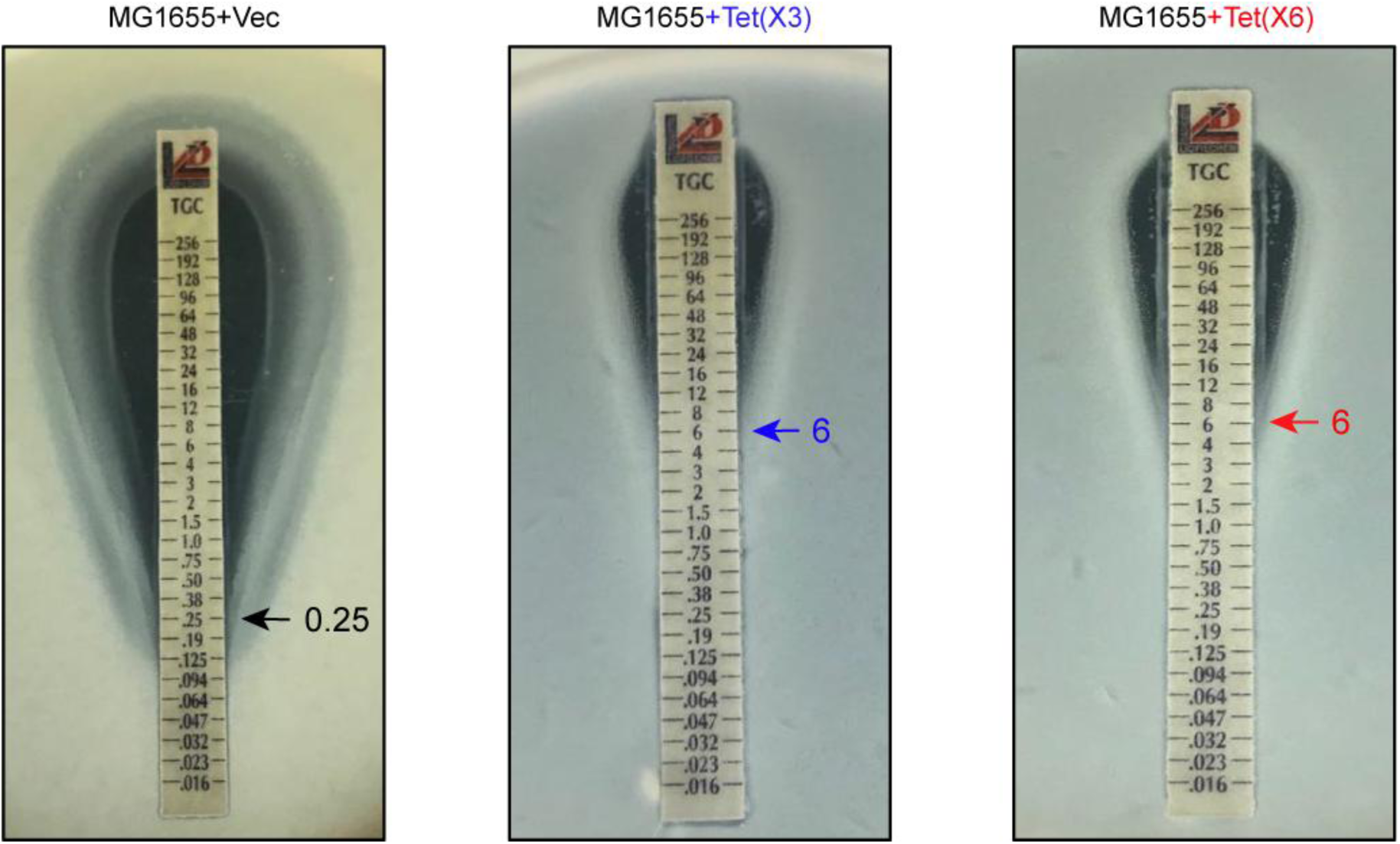
Use of E-test to evaluate phenotypic growth of MG1655 strains expressing either *tet*(X3) or *tet*(X6) Semi-solid medium was poured, which was supplemented with the MG1655 strains with the empty vector alone, or the plasmid-borne *tet*(X3) [and/or *tet*(X6)], accordingly. E-test strip is featuring with a gradient concentration of tigecycline. The cut-off value of tigecycline resistance is indicated with an arrow. A representative result of three independent experiments is given.

## Notes

### Competing Interest Statement

The authors have declared no competing interest.

### Summary of Updates

It is an improved version having multiple lines of text correction, image replacement, and new data!

## References

1. Li, J. et al. Colistin: the re-emerging antibiotic for multidrug-resistant Gram-negative bacterial infections. Lancet Infect Dis 6, 589–601 (2006).

2. Doan, T.L., Fung, H.B., Mehta, D. & Riska, P.F. Tigecycline: a glycylcycline antimicrobial agent. Clin Ther 28, 1079–1106 (2006).

3. Noskin, G.A. Tigecycline: a new glycylcycline for treatment of serious infections. Clin Infect Dis 41 Suppl 5, S303–14 (2005).

4. Feng, Y. Transferability of MCR-1/2 polymyxin resistance: Complex dissemination and genetic mechanism. ACS Infect Dis 4, 291–300 (2018).

5. Gao, R. et al. Dissemination and mechanism for the MCR-1 colistin resistance. PLoS Pathog 12, e1005957 (2016).

6. Liu, Y.Y. et al. Emergence of plasmid-mediated colistin resistance mechanism MCR-1 in animals and human beings in China: a microbiological and molecular biological study. Lancet Infect Dis 16, 161–8 (2016).

7. Wang, C. et al. Identification of novel mobile colistin resistance gene *mcr-10*. Emerg Microbes Infect 9, 508–516 (2020).

8. Sun, J., Zhang, H., Liu, Y.H. & Feng, Y. Towards understanding MCR-like colistin resistance. Trends Microbiol 26, 794–808 (2018).

9. Zhang, H., Srinivas, S., Xu, Y., Wei, W. & Feng, Y. Genetic and biochemical mechanisms for bacterial lipid A modifiers associated with polymyxin resistance. Trends Biochem Sci 44, 973–988 (2019).

10. Nguyen, F. et al. Tetracycline antibiotics and resistance mechanisms. Biol Chem 395, 559–75 (2014).

11. Li, W. et al. Mechanism of tetracycline resistance by ribosomal protection protein Tet(O). Nat Commun 4, 1477 (2013).

12. Park, J. et al. Plasticity, dynamics, and inhibition of emerging tetracycline resistance enzymes. Nat Chem Biol 13, 730–736 (2017).

13. Moore, I.F., Hughes, D.W. & Wright, G.D. Tigecycline is modified by the flavin-dependent monooxygenase TetX. Biochemistry 44, 11829–35 (2005).

14. Thaker, M., Spanogiannopoulos, P. & Wright, G.D. The tetracycline resistome. Cell Mol Life Sci 67, 419–31 (2010).

15. Yang, W. et al. TetX is a flavin-dependent monooxygenase conferring resistance to tetracycline antibiotics. J Biol Chem 279, 52346–52 (2004).

16. Volkers, G., Palm, G.J., Weiss, M.S., Wright, G.D. & Hinrichs, W. Structural basis for a new tetracycline resistance mechanism relying on the TetX monooxygenase. FEBS Lett 585, 1061–6 (2011).

17. Forsberg, K.J., Patel, S., Wencewicz, T.A. & Dantas, G. The tetracycline destructases: A novel family of tetracycline-inactivating enzymes. Chem Biol 22, 888–97 (2015).

18. He, T. et al. Emergence of plasmid-mediated high-level tigecycline resistance genes in animals and humans. Nat Microbiol 4, 1450–1456 (2019).

19. Sun, J. et al. Plasmid-encoded *tet*(X) genes that confer high-level tigecycline resistance in *Escherichia coli*. Nat Microbiol 4, 1457–1464 (2019).

20. Xu, Y., Liu, L., Sun, J. & Feng, Y. Limited distribution and mechanism of the TetX4 tetracycline resistance enzyme. Sci Bull (Beijing) 64, 1478–1481 (2019).

21. Guiney, D.G., Jr., Hasegawa, P. & Davis, C.E. Expression in *Escherichia coli* of cryptic tetracycline resistance genes from *Bacteroides* R plasmids. Plasmid 11, 248–52 (1984).

22. Speer, B.S., Bedzyk, L. & Salyers, A.A. Evidence that a novel tetracycline resistance gene found on two *Bacteroides* transposons encodes an NADP-requiring oxidoreductase. J Bacteriol 173, 176–83 (1991).

23. Thakare, R., Dasgupta, A. & Chopra, S. Eravacycline for the treatment of patients with bacterial infections. Drugs Today (Barc) 54, 245–254 (2018).

24. Forde, B.M. et al. Discovery of *mcr-1*-mediated colistin resistance in a highly virulent *Escherichia coli* lineage. mSphere 3, pii: e00486-18 (2018).

25. Xu, Y., Lin, J., Cui, T., Srinivas, S. & Feng, Y. Mechanistic insights into transferable polymyxin resistance among gut bacteria. J Biol Chem 293, 4350–4365 (2018).

26. Mao, J. et al. Antibiotic exposure elicits the emergence of colistin- and carbapenem-resistant *Escherichia coli* coharboring MCR-1 and NDM-5 in a patient. Virulence 9, 1001–1007 (2018).

27. Sun, J. et al. Co-transfer of *bla*NDM-5 and *mcr-1* by an IncX3-X4 hybrid plasmid in *Escherichia coli*. Nat Microbiol 1, 16176 (2016).

28. Li, X. et al. oriTfinder: a web-based tool for the identification of origin of transfers in DNA sequences of bacterial mobile genetic elements. Nucleic Acids Res 46, W229–W234 (2018).

29. Zhi, C., Lv, L., Yu, L.F., Doi, Y. & Liu, J.H. Dissemination of the *mcr-1* colistin resistance gene. Lancet Infect Dis 16, 292–3 (2016).

30. Gillings, M.R. Integrons: past, present, and future. Microbiol Mol Biol Rev 78, 257–77 (2014).

31. Zhang, A.N. et al. Conserved phylogenetic distribution and limited antibiotic resistance of class 1 integrons revealed by assessing the bacterial genome and plasmid collection. Microbiome 6, 130 (2018).

32. Chiou, C.S. & Jones, A.L. Nucleotide sequence analysis of a transposon (Tn5393) carrying streptomycin resistance genes in Erwinia amylovora and other gram-negative bacteria. J Bacteriol 175, 732–40 (1993).

33. Liu, D. et al. Identification of the novel tigecycline resistance gene *tet*(X6) and its variants in *Myroides, Acinetobacter* and *Proteus* of food animal origin. J Antimicrob Chemother 75, 1428–1431 (2020).

34. He, D. et al. A novel tigecycline resistance gene, *tet*(X6), on an SXT/R391 integrative and conjugative element in a *Proteus genomospecies* 6 isolate of retail meat origin. J Antimicrob Chemother 75, 1159–1164 (2020).

35. Peng, K., Li, R., He, T., Liu, Y. & Wang, Z. Characterization of a porcine *Proteus cibarius* strain co-harbouring *tet*(X6) and *cfr*. J Antimicrob Chemother 75, 1652–1654 (2020).

36. Wang, R. et al. The global distribution and spread of the mobilized colistin resistance gene *mcr-1*. Nat Commun 9, 1179 (2018).

37. Snesrud, E. et al. A Model for Transposition of the Colistin Resistance Gene mcr-1 by ISApl1. Antimicrob Agents Chemother 60, 6973–6976 (2016).

38. Yang, Q. et al. Balancing *mcr-1* expression and bacterial survival is a delicate equilibrium between essential cellular defence mechanisms. Nat Commun 8, 2054 (2017).

39. Zhang, H., Wei, W., Huang, M., Umar, Z. & Feng, Y. Definition of a family of nonmobile colistin resistance (NMCR-1) determinants suggests aquatic reservoirs for MCR-4. Adv Sci (Weinh) 6, 1900038 (2019).

40. Zhang, H. et al. Action and mechanism of the colistin resistance enzyme MCR-4. Commun Biol 2, 36 (2019).

41. Xu, Y. et al. Spread of MCR-3 colistin resistance in China: An epidemiological, genomic and mechanistic study. EBioMedicine 34, 139–157 (2018).

42. Zhang, H. et al. A genomic, evolutionary, and mechanistic study of MCR-5 action suggests functional unification across the MCR family of colistin resistance. Adv Sci (Weinh) 6, 1900034 (2019).

43. Wang, R.F. & Kushner, S.R. Construction of versatile low-copy-number vectors for cloning, sequencing and gene expression in *Escherichia coli*. Gene 100, 195–9 (1991).

44. Khlebnikov, A., Risa, O., Skaug, T., Carrier, T.A. & Keasling, J.D. Regulatable arabinose-inducible gene expression system with consistent control in all cells of a culture. J Bacteriol 182, 7029–34 (2000).

45. Song, M. et al. A broad-spectrum antibiotic adjuvant reverses multidrug-resistant Gram-negative pathogens. Nat Microbiol, https://doi.org/10.1038/s41564-020-0723-z (2020).

46. Wang, Q. et al. Complex dissemination of the diversified *mcr-1*-harbouring plasmids in *Escherichia coli* of different sequence types. Oncotarget 7, 82112–82122 (2016).

47. Stokes, J.M., Lopatkin, A.J., Lobritz, M.A. & Collins, J.J. Bacterial metabolism and antibiotic efficacy. Cell Metab 30, 251–259 (2019).

48. Lopatkin, A.J. et al. Bacterial metabolic state more accurately predicts antibiotic lethality than growth rate. Nat Microbiol 4, 2109–2117 (2019).

49. Belenky, P. et al. Bactericidal antibiotics induce toxic metabolic perturbations that lead to cellular damage. Cell Rep 13, 968–80 (2015).

50. Lobritz, M.A. et al. Antibiotic efficacy is linked to bacterial cellular respiration. Proc Natl Acad Sci U S A 112, 8173–80 (2015).

51. Kohanski, M.A., Dwyer, D.J., Hayete, B., Lawrence, C.A. & Collins, J.J. A common mechanism of cellular death induced by bactericidal antibiotics. Cell 130, 797–810 (2007).

52. Brynildsen, M.P., Winkler, J.A., Spina, C.S., MacDonald, I.C. & Collins, J.J. Potentiating antibacterial activity by predictably enhancing endogenous microbial ROS production. Nat Biotechnol 31, 160–5 (2013).

53. Dwyer, D.J. et al. Antibiotics induce redox-related physiological alterations as part of their lethality. Proc Natl Acad Sci U S A 111, E2100–9 (2014).

54. Sullivan, M.J., Petty, N.K. & Beatson, S.A. Easyfig: a genome comparison visualizer. Bioinformatics 27, 1009–10 (2011).

55. Caroff, M., Tacken, A. & Szabo, L. Detergent-accelerated hydrolysis of bacterial endotoxins and determination of the anomeric configuration of the glycosyl phosphate present in the “isolated lipid A” fragment of the *Bordetella pertussis* endotoxin. Carbohydr Res 175, 273–82 (1988).

56. Ye, H. et al. Diversified *mcr-1*-harbouring plasmid reservoirs confer resistance to colistin in human gut microbiota. mBio 7, e00177 (2016).

57. Balouiri, M., Sadiki, M. & Ibnsouda, S.K. Methods for *in vitro* evaluating antimicrobial activity: A review. J Pharm Anal 6, 71–79 (2016).

58. Walkiewicz, K. et al. Small changes in enzyme function can lead to surprisingly large fitness effects during adaptive evolution of antibiotic resistance. Proc Natl Acad Sci U S A 109, 21408–13 (2012).

59. Biasini, M. et al. SWISS-MODEL: modelling protein tertiary and quaternary structure using evolutionary information. Nucleic Acids Res 42, W252–8 (2014).

